# The Human Gut Cell Atlas v1.0: A Unified, Community-Built Reference of Cellular Diversity Across the Gut

**DOI:** 10.64898/2026.10.09.753994

**Authors:** Kyle Kimler, Christopher Lance, Balazs Bohar, Abdalla Ahmed, Ran Zhou, Raquel Bartolome-Casado, Victoria Gudino, Ricardo O. Ramirez Flores, Fredrik V. Junghus, Urs M. Morbe, Diana Domanska, Michaela F. Mueller, Madelyn R. Moy, Angela Sanzo-Machuca, Marton Olbei, Batuhan Cakir, Benjamin Weinert, Benjamin M. Frey, Michael Glinka, Tobi Alegbe, Duy Pham, Joshua de Sousa Casal, Alexander V. Predeus, Alexandre Denadai-Souza, Francesca Perrone, Hyun Chang Lee, Sushma Chowdary Maddipatla, David Rogers, Mary K. Futey, Arsenios Chatzigeorgiou, Ida Zucchi, Jasmine Belfiore, Lucia Robson, Agne Antanaviciute, Daniel B. Rainbow, Tatiana A. Karakasheva, Richard Baldock, Derek Houghton, Vladimir A. Shitov, Chenchen Zhu, Chloe Hyun-Jung Lee, Elva Maria Novoa-del-Toro, Sara Kimler, Anna Hupalowska, Guy E. Boeckxstaens, Eugene C. Butcher, Lori A. Coburn, Shahida Din, Jellert T. Gaublomme, Narayanan Kasthuri, Bartha M. Knoppers, Dermot P. B. McGovern, Jeffrey R. Moffitt, Andreas E. Moor, Peng Qiu, Keith T. Wilson, the Helmsley Gut Cell Atlas Crohn’s Disease Consortium, Candace M. Cham, Eugene B. Chang, Can Ergen, Donna L. Farber, Mennatallah Ghouraba, Jason Koval, Christopher J. Lengner, Michal Mark, David T. Rubin, Peter A. Sims, Steven B. Wells, Bingqing Xie, Nir Yosef, Yusen Zhou, David Fawkner-Corbett, Thomas M. Fenton, Scott T. Magness, Shalev Itzkovitz, Monika Krzak, Rebecca E McIntyre, Adrienne Luoma, Gregory J. Wickham, Joseph Burclaff, Shengbao Suo, Liza Konnikova, Keith Alexander Breau, Weihong Gu, Espen S Baekkevold, the Human Cell Atlas Gut Bionetwork, D Leland Taylor, Joanne L. Jones, John W. Hickey, Carl A Anderson, Tim Raine, Julio Saez-Rodriguez, Sebastian Pott, Irene Papatheodorou, David J Adams, Alison Simmons, Albert Burger, Tamas Korcsmaros, Mark J Arends, Anindita Basu, Kathryn E. Hamilton, William W. Agace, David Osumi-Sutherland, Kylie R. James, Amanda J. Oliver, Rasa Elmentaite, Michael P. Snyder, Sarah A. Teichmann, John Randell, Azucena Salas, Fabian J. Theis, Malte D. Luecken, Daniel Kotlarz, Matthias Zilbauer, Jose Ordovas-Montanes

## Abstract

Single-cell RNA-sequencing (scRNA-seq) studies have transformed our understanding of cellular diversity in the human intestine, but differences in anatomical sampling, experimental processing and cell-type nomenclature limit comparison across datasets. Here, we present the Human Gut Cell Atlas (HGCA) V1, a community-curated reference of 944,390 healthy and disease-adjacent intestinal cells from 502 samples, 265 donors and 24 landmark studies, linked to detailed anatomical, clinical, and technical metadata. As this metadata is of critical importance to human biology, we developed MetaManager, an open source data management framework for standardized metadata collection and tracking. A five-level consensus taxonomy resolves 94 homeostatic cell identities and enables transferrable, high-resolution annotation of new gut datasets. By assembling cell identity, sampling metadata, and independent spatial validation, intestinal segment and radial tissue depth were revealed to systematically shape cellular composition and transcriptional state, enabling inference of the presence of sampling-dependent tissue niches, including lymphoid follicles. Reference mapping accurately and rapidly recovered cohort-level Crohn’s disease and treatment signatures without *de novo* annotation, while mapping human intestinal organoids quantified cell-type-specific identity, regional specification, maturation, and transcriptional fidelity to native tissue. The HGCA V1 provides a standardized cellular, anatomical coordinate, and metadata system for reproducible, high-quality analysis of intestinal biology across healthy tissue, disease-associated remodelling, and experimental models, providing a continuously evolving community reference for cumulative discovery in human intestinal biology.

## Introduction

The human gastrointestinal tract is one of the most anatomically and cellularly complex organ systems in the body. Along the intestines’ longitudinal axis, specialized regions from the duodenum to the rectum perform distinct physiological functions in nutrient absorption, epithelial renewal, barrier maintenance, immune surveillance and host-microbiome interactions^1^. Epithelial, immune, stromal, vascular, neuronal and muscular cell populations are organized into specialized microenvironments spanning crypt-villus units, mucosal and submucosal compartments, spatially discrete lymphoid follicles and mesenchymal niches. Cell composition and transcriptional programs therefore vary substantially with both axial (i.e. length) and radial (i.e. depth) position.

This complexity is directly reflected in human disease. Gastrointestinal diseases represent a major global health burden, affecting hundreds of millions of individuals worldwide. Notably, they exhibit remarkably heterogeneous pathological remodeling, selectively affecting distinct intestinal regions, tissue compartments and cellular populations. Crohn’s disease (CD) predominantly affects the terminal ileum and the nearby proximal colon, and frequently extends through the bowel wall^2^, whereas ulcerative colitis is largely confined to the colonic mucosa^3^ and celiac disease targets the proximal small intestine^4^. Consequently, distinguishing physiological regional specialization from disease-associated remodeling requires a comprehensive reference of healthy intestinal cellular diversity across anatomical locations.

Single-cell RNA-sequencing (scRNA-seq) has uncovered previously unrecognized intestinal populations and tissue programs, including BEST4-positive enterocytes^5,6^, villus epithelial zonation^7^, distinct crypt microenvironments^8^, pyloric and Brunner’s gland-like metaplastic epithelial cells^9^, and fibroblast-immune niches associated with intestinal fibrosis^10,11^. Collaborative initiatives have begun to consolidate these data: PanGI^9^ demonstrated the feasibility and utility of cross-study integration, HuBMAP^12,72^ established anatomically anchored tissue references, and the Helmsley Gut Cell Atlas contributed a common coordinate framework^13^ and clinically structured cohorts. Despite these important advances, existing studies and reference atlases still differ substantially in anatomical sampling, tissue collection, metadata standards, analytical workflows and cell-type nomenclature, resulting in inconsistent annotation of equivalent cell populations, incomplete separation of biological and technical variation, and a risk of unstable interpretation of disease signatures. The limited availability of healthy gut tissue further constrains the scale and population diversity in current references, and the lack of standardized metadata and a versioned mechanism for continuous revision and expansion will lead to a gradual loss of atlas relevance as knowledge on cell identities accumulates and new data or modalities are introduced.

To overcome these limitations, the three single-cell gut consortia came together with the wider gastrointestinal research community to form the Human Cell Atlas Gut Bionetwork to establish the Human Gut Cell Atlas (HGCA), a community-built and continuously evolving reference. The HGCA v1.0 comprises 944,390 cells from 502 healthy or disease-adjacent normal tissue samples obtained from 265 donors across 24 landmark studies (**Extended Data Table 1**)^5,6,10,14–36^. Using two international expert annotation jamborees, we developed a five-level hierarchical taxonomy of intestinal cell identities that is transparently curated through the Cell Annotation Platform (CAP), where supporting evidence, expert review and consensus decisions are permanently documented and continuously refined as new biological knowledge emerges. We demonstrate accurate transfer of this framework to a Crohn’s disease cohort and human intestinal organoids, enabling the rapid and robust detection of disease signatures and assessment of *in vitro* fidelity to native tissue. These analyses establish the HGCA as a standardized foundation for interpreting intestinal homeostasis, disease-associated remodeling, and experimental model systems, while providing a community framework that can continuously evolve with new datasets, technologies and biological knowledge.

## Results

### Generating a high-quality reference atlas of the human gut

The HGCA v1.0 assembles 944,390 single-cell profiles from 502 samples and 265 donors covering duodenum, jejunum, ileum and colorectum (hereafter colon) across the human lifespan (**Fig 1a-b**). We integrated 24 datasets from 20 countries after reviewing 60 candidate studies (representing over 1000 samples and 6.2 million cells) using predefined inclusion/exclusion criteria based on healthy reference tissue, droplet based single cell technologies, metadata completeness and expert representation of intestinal regions (**Supplementary Fig. 1**). Datasets focused on the stomach and esophagus were excluded due to their distinctive epithelial composition and will be incorporated in future efforts. The final atlas contains 613,002 cells from the small intestine (65%) and 289,028 from the colon (31%). Most cells derive from healthy mucosa (∼78%), complemented by disease-adjacent tissue (∼22%) to broaden study inclusion. The atlas includes both endoscopic biopsies (194 donors; 73.2%) and surgical resections (71 donors; 26.8%), and consists exclusively of scRNA-seq 10X Genomics transcriptomes (10X 3’ v3: 56.9%, 10X 3’ v2: 26.3%, 10X 5’ v1: 10.1%, 10X 5’ v2: 4.5%, 10X 5’ v3: 2.2%). The current release is enriched for adult donors of predominantly European ancestries, reflecting currently available data. The atlas spans ages from birth to 89 years of age (∼10% of cells pediatric/adolescent), including balanced representation of sex and capturing the major intestinal compartments (61% lymphoid and 29% epithelial populations). Together, HGCA v1.0 provides a comprehensive reference for adult intestinal mucosa while defining a scalable framework for continued expansion across populations, anatomical regions and molecular modalities.

**Figure 1.**
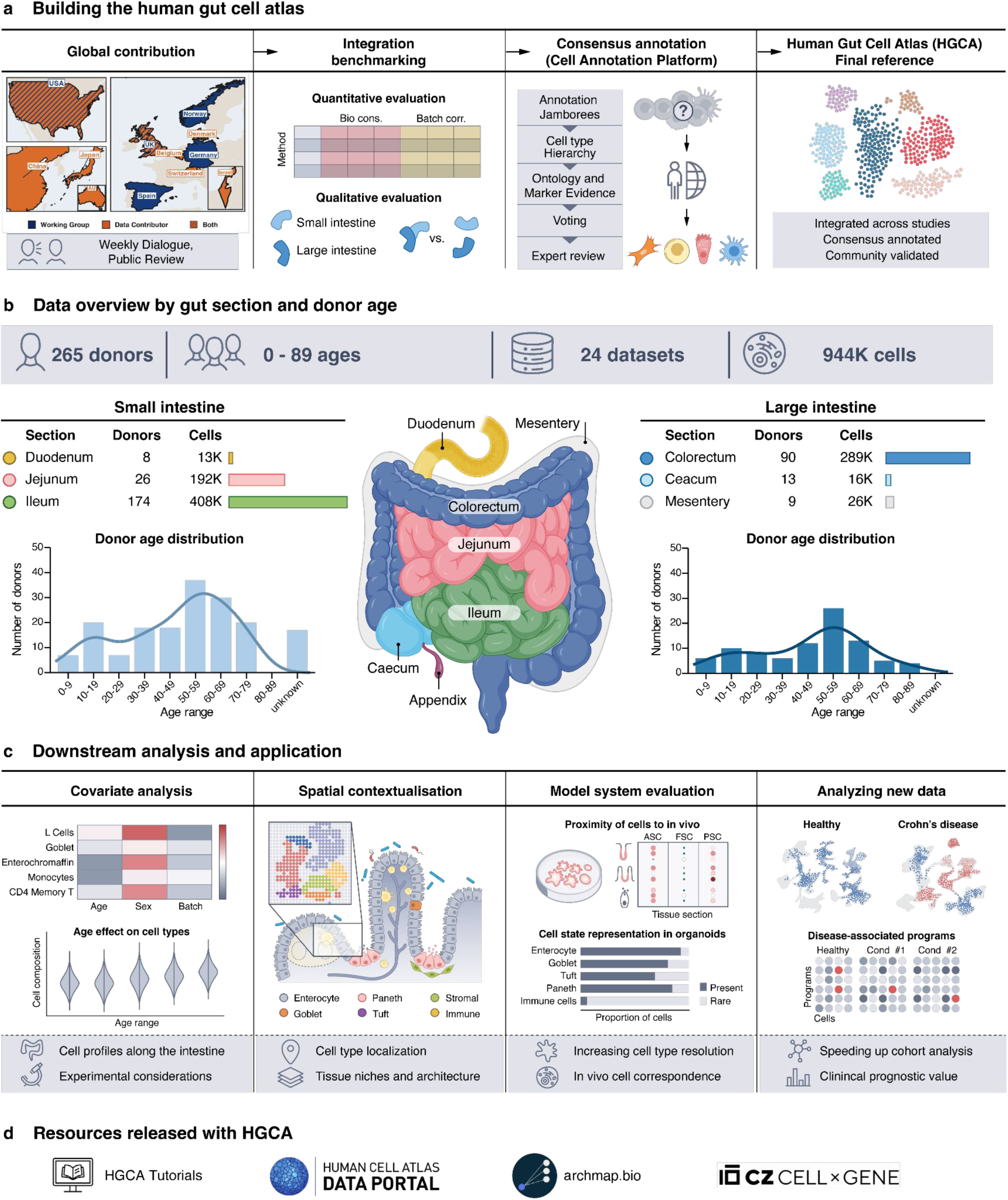
The Human Gut Cell Atlas (HGCA) overview and cohort composition. a) HGCA construction via global contributions, integration benchmarking, followed by consensus annotation across the gut bionetworks using the Cell Annotation Platform (CAP). b) The gut cell atlas by the numbers, including 265 donors spanning 0-89 years of age, 24 datasets, and 944,390 cells of the small and large intestines c) Downstream analysis and use cases of the HGCA demonstrated in the paper, including (i) covariates analysis in healthy data, (ii) spatial contextualisation, (iii) intestinal gut organoids in vivo correspondence, and (iv) mapping and rapidly analyzing disease data. d) Iconography of applications and resources provided with the atlas

The construction of the HGCA was guided by three principles: metadata harmonization, benchmarked integration, and community consensus. We considered metadata an integral component of the reference because biological interpretation depends not only on transcriptomes but also on a long list of experimental and biological factors, including tissue origin, sampling strategy, processing, and the many details of clinical context. Defining a healthy gut baseline is complicated because control tissues are obtained from diverse sources, including endoscopic screenings, surgery, post-mortem donation, or disease-adjacent sampling. Samples destined for single-cell analysis may also then be pinched, sliced, heated, chilled, frozen, or bathed in chemicals and enzymes before their transcriptional profile is measured. These sampling differences affect cellular profiles, but are not documented in a standardized way across studies **(Supplementary Fig. 2)**. Consequently, the quality and value of a reference atlas depends as much on the completeness, provenance and traceability of its metadata and annotations as on its cellular scale. To standardize metadata collection, we built a code library^37^, MetaManager, an open source data management framework that has been incorporated into the HCA website (https://tracker.humancellatlas.org) for coordinated metadata collection and live metadata tracking for atlas builders and contributors. Standardized metadata templates and public dashboards provide instant feedback on completeness, supporting international curatorial collaboration.

To maximize biological fidelity, we systematically benchmarked integration strategies using scAtlasTb^38^, a benchmarking and atlas building toolbox, to select the best method for removing batch effects while preserving biological information. For the HGCA, the dataset of origin was the greatest contributor to technical variance in expression profiles followed by the 10X assay used. Within-lineage (i.e. separating epithelial, stromal, lymphoid and myeloid) scVI integration over donor ID performed best based on quantitative metrics^39^ and manual identification of nuanced cellular identities (**Supplementary Fig. 3**).

Community consensus was established through a versioned, evidence-backed annotation process through the Cell Annotation Platform (CAP). Two annotation jamborees brought together experts spanning developmental, spatial, and clinical gut biology to evaluate candidate identities using marker expression, anatomical localization, cross-dataset reproducibility, and prior literature evidence. CAP records the evidence, reviews, expert discussion and consensus decision underlying every annotation, and remains open after publication for community-driven refinements as new datasets emerge (**Fig. 1c**). Together with interactive visualization, accessible reference mapping methods, and programmatic access (**Fig. 1d**), this establishes the HGCA as a continuously evolving community reference for the human intestine.

### Consensus annotation defines standardized intestinal cell identities

Lineage-specific integration of epithelial, lymphoid, myeloid, and stromal populations followed by iterative clustering identified 574 candidate populations for expert annotation. Together with the Human Cell Atlas Gut Bionetwork, we established a hierarchical taxonomy comprising 94 consensus intestinal cell types organized across five annotation levels and linked each identity to canonical markers, Cell Ontology terms, and literature evidence (**Fig. 2a,b, Extended Data Table 2**). Community curation through the CAP generated a graphical taxonomy documenting annotation evidence, comparison with original author labels, expert consensus, and dataset representation for every cell type (**Fig. 2c**). Thirty-two of the consensus 94 cell types each represented less than 0.1% of all cells (“Rare”), while quality control metrics and dataset diversity demonstrate the robustness of the identified cell types, demonstrating that rare populations can be reproducibly resolved across independently generated datasets.

**Figure 2.**
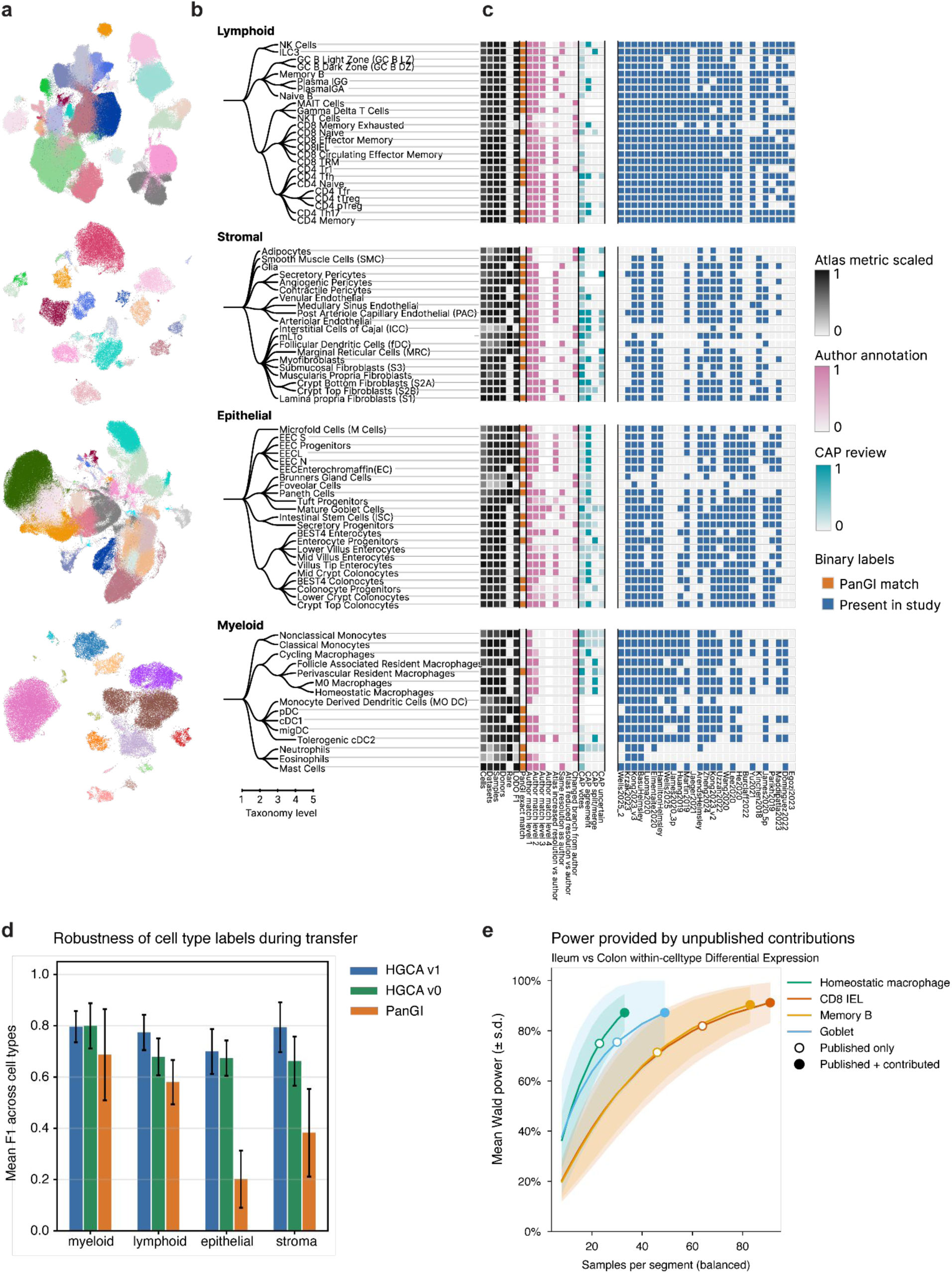
Evidence based taxonomy of healthy human gut celltype. a) Uniform Manifold Approximation and Projection of integrated cell embeddings per cell type lineage colored by cell type. b) Five-level HGCA v1 celltype hierarchies for epithelial, lymphoid, myeloid, and stromal lineages. Nodes are consensus identities and edges are curated parent-child relationships. c) Aligned heat maps summarize, for each high-resolution celltype, atlas support (grey), exact presence in PanGI healthy atlas (orange), hierarchical concordance with original author annotations mapped through their closest HGCA celltype (pink), CAP review metrics (teal), and binary presence across datasets (blue). d) Leave-one-dataset-out label-transfer benchmark comparing the PanGI healthy reference, HGCA v0 annotations before Cell Annotation Platform review, and HGCA v1 annotations after community review. Bars show mean macro F1 across held-out datasets for each lineage, error bars show variation across held-out datasets. e) Power analysis for differential expression (DE) between ileum and colon, showing increase in DE genes found once unpublished datasets are included. Samples included in the comparison are balanced according to the minimum number of them in either ileum or colon. DESeq2 formula included removal of dataset-specific batch effect.

To quantify the impact of community consensus, we compared HGCA annotations with the nearest HGCA equivalents of the original author annotations. Labels differed for 738,015 of 944,390 compared cells (78.14%). In the HGCA, 565,331 cells (59.86%) received a higher-resolution label than in the original study (refinement of cellular identity), while 162,616 cells (17.22%) were re-annotated to a different taxonomy branch (initial misannotation), 10,068 cells (1.07%) received a coarser label (potential contentious original author labels), and 206,487 (21.86%) retained the original label (**Fig. 2c** pink). The most frequent reassignment between taxonomy branches was from CD4 to CD8 T cells and vice versa, highlighting appreciated challenges of resolving these populations using transcriptomic data. CAP metrics further documented the degree of expert discussion underlying each annotation, indicating that lymphoid, stromal, and dendritic cell populations received fewer instances of feedback than the rest (**Fig. 2c** teal; **Supplementary Fig. 4**). Consensus review resolved several nomenclature inconsistencies, including the most-upvoted piece of feedback, that “Secretory Goblet Cells” should be renamed “Goblet Cells”. A second highly up-voted change was to split the “M2 Macrophages” category into multiple anatomically and functionally defined tissue-resident macrophage subtypes.

The HGCA v1 substantially expands the resolution and consistency of intestinal cell identities by resolving numerous rare or previously inconsistently annotated cell populations across studies. Examples include eosinophils, medullary sinus endothelial cells, T follicular regulatory cells, follicle-associated resident macrophages, and tuft progenitors. High-resolution epithelial annotation distinguished BEST4 enterocytes and BEST4 colonocytes as zonation-resolved entities and reidentified PanGI’s INFLARE cells as subclusters of Brunner’s gland and foveolar cells in disease-adjacent control samples (**Supplementary Fig. 5**). Absorptive epithelial lineages were resolved into five discrete cell states (TA, segment-committed progenitors, early, mid, and late) spanning the crypt-surface axis in the colon and the crypt-villus axis of the small intestine, whereas enteroendocrine cells were subdivided into five peptide-defined populations (EEC L, EEC N, Enterochromaffin, EEC S, and EEC progenitors).

Stromal annotation organized fibroblasts according to their spatial distribution and anatomical localization within the intestinal wall, distinguishing lamina propria populations, crypt top and bottom fibroblasts, submucosal fibroblasts, and muscularis propria fibroblasts, as set out by previous scRNAseq studies^25^. Myofibroblasts, reticular fibroblasts, mLTo cells, and ICCs share an overarching level 2 annotation with these fibroblasts which separates them from pericytes, endothelial cells, glia, smooth muscle cells, and adipocytes. Pericytes were subdivided into secretory, angiogenic, and contractile populations, consistent with functional specialization and differential microlocalization^19^. Within the lymphoid compartment, conventional and unconventional T cells, B cells and innate lymphoid cells were further subdivided into naïve, memory and tissue-resident populations, including intraepithelial CD8 T cells (CD8 IEL)^22^, tissue-resident memory CD8 T cells (CD8 TRM)^40^, and germinal center B cells of the light and dark zones (GC B LZ; GC B DZ)^41^. Regulatory T cells (Tregs) were further subdivided into Helios+/- groups which were named thymic (tTreg) and peripheral (pTreg)^42^. Similarly, the myeloid lineage distinguished dendritic cells, granulocytes, monocytes, and macrophages, and anatomically specialized tissue-resident macrophage populations, separating M0 macrophages (lacking CD209 (MRC1), FOLR2, and CD163L1) from cytokine-conditioned homeostatic macrophages on the basis of conserved transcriptional programs^43^.

The practical value of a reference atlas depends on how reliably its annotations generalize to new datasets. To evaluate how HGCA atlas design principles compare to existing meta-atlases and single-cohort studies, we tested cell-level label transfer using leave-one-dataset-out (LODO) cross-validation (Methods). Relative to both HGCA v0 and PanGI, HGCA v1 refined annotations for several healthy intestinal cell types, including Tregs and macrophage subtypes (**Fig. 2a** orange). These refinements translated into substantially improved transfer performance, increasing mean cell-type F1 from 0.68 to 0.78 in lymphoid, from 0.67 to 0.7 in epithelial, and from 0.66 to 0.79 in stromal lineages between HGCA v0 and v1. Under the same scANVI evaluation, HGCA v1 also exceeded PanGI across all four lineages, with the largest improvement within the epithelial lineage (ΔF1 ≈ 0.5; **Fig. 2d**). Cell-type-specific transfer performance is summarized in **Fig. 2c** (LODO F1 black), demonstrating that community consensus substantially improves the robustness and transferability of intestinal cell identity annotation.

Maintaining a living, versioned community atlas requires continuous integration of new datasets and transparent, trusted community governance. HCA collaboration infrastructure and metadata tracking provided this visibility into available datasets, contribution, and remaining biological gaps, facilitating inclusion of unpublished datasets and accelerating atlas expansion. The inclusion of these additional datasets substantially improved statistical power to resolve region-specific biology, exemplified by differential expression between the ileum and colon across multiple cellular lineages (**Fig. 2e; Supplementary Fig. 6**). Community contributions demonstrate the HGCA’s ability to evolve as a continuously updated community reference, progressively improving the resolution, consistency, and transferability of a shared intestinal cell nomenclature as new datasets and biological knowledge emerge.

### HGCA metadata disentangle anatomical and experimental determinants of intestinal single-cell datasets

A central challenge in interpreting single-cell atlases is distinguishing biological variation from differences introduced by anatomical variation and experimental design. Using the detailed anatomical, clinical, and technical metadata of the HGCA, we quantified variation in cell-type composition and transcriptional state across intestinal segment, radial tissue layer, donor characteristics and experimental protocols. By encoding sample collection method and radial tissue fractionation in metadata, samples could be positioned along two major spatial axes of the gut, radial tissue depth and longitudinal intestinal segment, consistent with emerging common coordinate frameworks of the gut^13^. To quantify the distribution of cell types along these axes, we calculated their relative within-lineage centered log-ratio (CLR) transformed compositions ( **Fig. 3a**).

**Figure 3.**
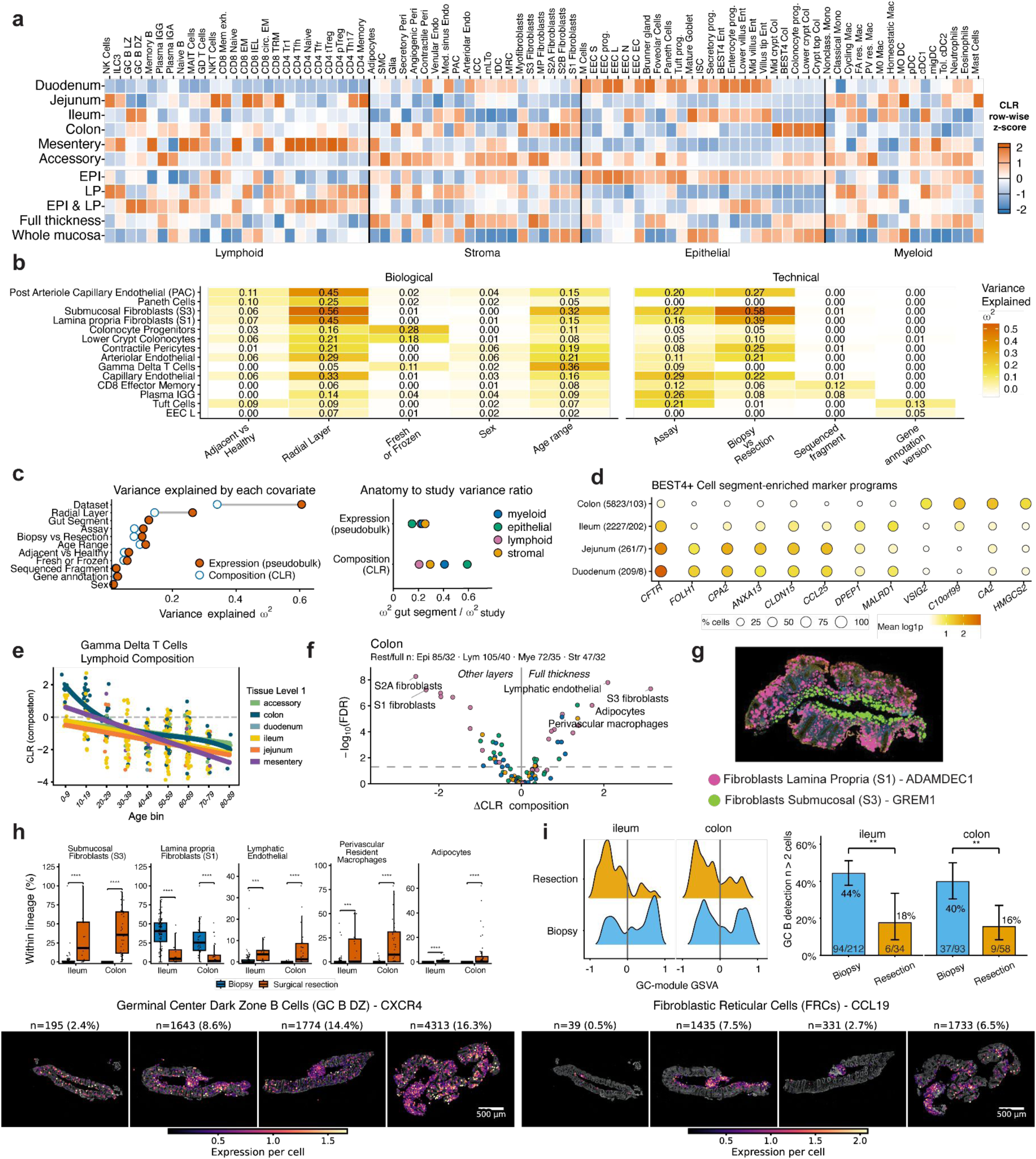
Anatomical and technical context changes the cellular view of the gut. a) Relative compositions (row-wise z-score; CLR transformed) of cell types across major segments and radial depth sampling across the gut. b) Heat map of the two cell types with the largest composition effect size for each covariate, grouped into biological and technical covariates. Colour denotes variance explained ωZ c) Variance in cell type composition and pseudobulk expression associated with each covariate. The gut segment to study ratio is plotted by lineage for centred log ratio (CLR) composition and pseudobulk expression. d) Segment enriched marker programs in BEST4 enterocytes and BEST4 colonocytes, cell numbers / samples represented on Y axis. e) gamma-delta T cell population reduces with age across gut segments. f) Differential cell type composition between full thickness specimens and samples containing one or more superficial radial layers in ileum and colon. g) 10X Xenium spatial assay of colon showing localization of S1 and S3 fibroblasts by depth. h) Percent capture of depth-localized cell types by resection or biopsy. i) (left) Sample-level GSVA score distributions for GC B-associated gene modules (right) Fraction of samples with >2 GC B cells among biopsies vs resections. j) Xenium spatial transcriptomics images of 4 colon biopsies. Dots represent centroids of cells, colored by normalized expression of the target gene.

When designing a new study, choices about microanatomical location, experimental protocols, and specimen conditions can determine which cell populations are recoverable and how comparable samples will be across cohorts. The HGCA provides a quantitative catalog of these associations and their limits of interpretability (**Fig. 3b-c**). To understand how experimental protocol influences dataset composition, we examined these covariates at two levels: cell-type composition, asking how the relative abundance of populations within a lineage varies across samples (e.g., subepithelial versus submucosal fibroblasts), and within-cell-type gene expression, asking how the transcriptional state of the same cell identity varies across samples (e.g., BEST4+ absorptive cells across intestinal segments). For each analysis, we summarized the strength of association with individual anatomical, biological, and technical covariates using an effect-size metric estimating the variance associated with a covariate (ω^2^; principal-component regression). Study of origin remained the strongest determinant of transcriptional variation across all lineages [composition 0.309-0.396; expression 0.525-0.699], reflecting differences in sample processing and experimental design. Among anatomical coordinates, radial tissue depth explained more variation than axial gut segment for both cellular composition [lymphoid: 0.107, myeloid: 0.123, epithelial: 0.167, stroma: 0.180] and transcriptional state [myeloid: 0.204, lymphoid: 0.212, stroma: 0.246, epithelial: 0.410], whereas axial gut segment (duodenum, jejunum, ileum, colorectum) explained less variance in both analyses [composition 0.082-0.187; expression 0.104-0.152]. Thus, among anatomical and sampling variables, radial depth is a major determinant of the cellular landscape recovered by existing intestinal scRNA-seq datasets.

Under the standardized HGCA annotations, composition retained biological structure more strongly relative to study-associated variation than within-cell-type expression. Accordingly, the ratio of variance explained by gut-segment to variance explained by study of origin (ω^2^) was consistently higher for composition than for pseudobulk expression (**Fig. 3c**, right), indicating that cellular composition retains biological signal while being less susceptible to study-specific effects than gene expression. Because anatomical, clinical, and technical variables are frequently interdependent, we quantified pairwise covariate dependence using Theil’s U and provide the resulting matrix as a guide to interpretation (**Supplementary Fig. 7**). For example, study of origin strongly predicted whether tissue was obtained by biopsy or resection (Theil’s U = 0.98), reflecting study-specific sampling strategies, while age range showed only weak dependence on disease status (U = 0.25). Quantifying these relationships distinguishes largely independent covariates from those describing overlapping biological or experimental features, enabling more robust interpretation of downstream associations.

Radial sampling illustrates the value of this metadata framework. Endoscopic biopsies predominantly sample the mucosa, whereas surgical resections may include submucosa, muscularis, and serosa. The sampling method does not uniquely determine the tissue ultimately sequenced, however, as biopsies and resections may be fractionated by enzymatic digestion, with only one or several radial layers carried forward for sequencing. Collection method and radial layer therefore encode overlapping but distinct information (U = 0.39), with both showing strong effects on cellular composition and expression. Their high ω^2^ points to the importance of careful consideration of these protocols. At the compositional level, radial sampling primarily affected the recovery of spatially restricted populations, including deep fibroblasts and associated stromal states. Radial depth also influenced transcriptional states within shared cell identities. For example, enteric glia (ω^2^ = 0.20) showed increased expression of RELN in lamina propria (LP) samples (log_2_FC 3.67; p=0.034), a marker of type III glia localized to the intestinal mucosa, whereas glia from full-thickness samples exhibited higher APOD expression (log2FC 2.49; p=0.019), characteristic of extraganglionic type IV glia in the muscularis, together with ENTPD2 (log2FC 3.02; p=0.083), and SFRP5 (log2FC 2.99; p=0.061).^44^ Thus, radial sampling determines not only which cell populations are recovered but also which spatially specialized states are captured within the same cell identity. The HCGA can therefore inform how tissue collection and fractionation may shape both cellular composition and transcriptional state.

To extend this framework across anatomical, clinical, and technical metadata, we ranked cell-type-specific composition effects for each covariate (**Fig. 3b**). Among the largest effects were radial layer for submucosal fibroblasts (ω^2^ 0.56) and post-arteriole capillary endothelial cells (ω^2^ 0.45), age for γδ T cells (ω^2^ 0.36), and tissue preservation method for colonocyte progenitors (ω^2^ 0.29). Together with the covariate-dependence matrix, these rankings provide practical guidance for identifying experimentally relevant differences in cell recovery (**Extended Data Table 3**). For example, freezing substantially reduced goblet cell recovery (ΔCLR = -1.85, Wilcoxon rank-sum test FDR = 1.1×10^-6^), whereas perivascular resident macrophages were enriched in full-thickness ileal samples (ΔCLR = 2.48, FDR = 6.1×10-5), and γδT cells progressively decreased with age across all gut segments (Spearman r = -0.50, n = 290; **Fig. 3e**). Along the axial direction of the gut, small intestinal segments were enriched for CD8 T resident memory cells, CD8 intraepithelial lymphocytes, NKT cells, and transiently amplifying (TA) epithelial cells, while cDC1s, goblet cells, and perivascular resident macrophages were preferentially recovered from the colon (**Supplementary Fig. 8**).

The enrichment of perivascular macrophages in deep tissue reflected a broader compositional shift with sampling depth. In the colon, full-thickness and more superficial preparations differed across 47 cell types, with 22 enriched in full thickness and 25 in the individual or superficial layers (**Fig. 3f**). Across both ileum and colon, full-thickness samples were enriched for submucosal fibroblasts (S3), lymphatic endothelial cells, perivascular resident macrophages, and adipocytes, with corresponding depletion of lamina propria (S1) and crypt bottom (S2A) fibroblasts. The shared S3-lymphatic-perivascular signal suggests that increasing sampling depth preferentially captures a vascularized crypt-support region spanning the deep mucosa and submucosa, whereas adipocytes and muscular propria-associated fibroblasts indicate more limited additional recovery of outer bowel wall compartments^8,45,46^. Together, these findings demonstrate that anatomical location and sampling depth are major determinants of the cellular landscape recovered by intestinal single-cell transcriptomics and highlight the value of harmonized metadata for biological interpretation, experimental design and cross-study comparison.

### Spatial transcriptomics validates anatomical inference from HGCA metadata

Although harmonized metadata enables broad anatomical placement of scRNA-seq samples, dissociated transcriptomes lack the spatial context required to directly localize cell populations within the intestinal wall. We used independent spatial transcriptomic datasets, including healthy 10X Visium sections and newly generated Xenium profiles (Prime 5K Human Pan Tissue & Pathways Panel) to validate anatomical inferences derived from the HGCA. Xenium data displayed patterns associated with the shallow mucosal layer (ADAMDEC1, S1 fibroblasts) and the deeper vasculature best captured by full thickness scRNAseq samples (GREM1, S3 fibroblasts) (Fig. 3g). Cell2Location HGCA label marker scoring of 10X Visium healthy sections from across the gut recovered several higher-order tissue niches (**Supplementary Fig. 9**). Notably, perivascular macrophages and S3 fibroblast consistently co-localized within a shared niche (factor 4) associated with mast cells rather than lymphatics, while lymphatic endothelial cells localized to a distinct vascular compartment with only limited overlap. This niche (Factor 4) localized predominantly to the deep mucosa of a healthy 10X Visium colon sample, independently corroborating the depth-associated compositional differences identified in the HGCA (**Fig. 3h**). Spatial mapping also highlighted regions where current reference data remain incomplete. For example, adipocytes clustered with other rare cell populations, including neutrophils and monocyte-derived dendritic cells, suggesting that the outer wall niche may yet not be sufficiently captured by existing full thickness scRNA-seq datasets. In contrast, lymphoid follicle niche patterning was robustly identified across both 10X Visium (**Supplementary Fig. 9**; factor 8) and Xenium (Fig. 3j) datasets through co-localization/expression of CXCR4, a marker of Germinal Center Dark Zone B cells (GC B DZ) and CCL19, which marks Fibroblastic Reticular Cells (FRCs). Together, these independent spatial datasets validate the anatomical relationships inferred from HGCA metadata, demonstrating that harmonized metadata capture biologically meaningful higher-order tissue organization extending beyond individual cell identities.

### Segment-specific function distinguishes epithelial identities and modulates immune cell states

The relationship between cell identity and anatomical position differs fundamentally across intestinal segments, so a cell type taxonomy must treat each segment accordingly. An enterocyte and colonocyte share a lineage and an absorptive role but perform fundamentally different physiological functions, and the HGCA therefore assigns them as separate identities. Testing small intestine (SI) against large intestine (LI) function within absorptive cells consequently required collapsing these identities into a common Absorptive Epithelial node then split up by maturity. In contrast, most lymphoid and myeloid cells recur throughout the intestine, where they adopt segment-specific transcriptional programs while maintaining a shared cellular identity. To distinguish intrinsic cell identity from regional specialization, we compared Gene Ontology biological process enrichment between SI and LI for each cell type with sufficient donor support (**Supplementary Fig. 10**). Regional differences were most pronounced within the epithelium. SI and colonic absorptive epithelial cells exhibited distinct digestive, metabolic and absorptive programs, consistent with their classification as separate epithelial identities, while SI epithelium additionally showed enrichment of antigen presentation pathways. In contrast, shared immune populations largely retained the same cellular identity across intestinal segments but displayed region-specific functional adaptation. For example, LI myeloid and lymphoid cells were enriched for differentiation and activation programs such as antigen receptor-mediated signaling pathways in DC and B cell subtypes. Together, these patterns indicate that the immune interface is organized differently along the gut, with epithelial antigen presentation predominating in the SI and resident immune cell activation predominating in the colon.

Segment-dependent transcriptional programs also refined the HGCA taxonomy. For example, BEST4 labelled cells overall had a segment-specific expression axis determined by proximal to distal localization, recalling single-study cross-tissue atlases that found the same^19^. CFTR marked proximal BEST4 enterocytes and declined toward the distal gut, whereas colonic BEST4 were marked by VSIG2, C10orf99 (GPR15L), CA2, and HMGCS2 (**Fig. 3d**). Similarly, circulating and tissue resident T cells retained shared cellular identities while expressing established segment-specific homing receptors, with CCR9 enriched in the SI and GPR15 in the colon, recapitulating established segmental markers of intestinal T cells^47^, but as these cells are dynamic and circulate, they were kept under a general nomenclature. Among shared stromal populations, lamina propria fibroblasts showed the strongest segment-related transcriptional differences (including colon-biased POSTN), while venular MADCAM1/ACKR1 addressin programs were largely conserved across segments rather than region-restricted^48^. Collectively, these analyses distinguish cellular identities that are intrinsically linked to anatomical location from those whose regional specialization is better represented as context-dependent transcriptional programs. This provides a biological rationale for the HGCA five-level hierarchical taxonomy which separates stable cell identity from anatomical context while allowing both to be refined as new evidence emerges on tissue localization and the persistence of shared cellular programs across intestinal regions.

### Inferring lymphoid follicle capture and niche-specific cellular communication with the HGCA

In gut tissue, tertiary lymphoid structures (TLS) and lymphoid follicles are spatially distinct structures dispersed throughout the tissue in large aggregates that may be completely missed by tissue sampling. HuBMAP previously defined an inner follicle niche corresponding to the germinal center and characterized by enrichment of B cell populations^8^. To test whether this niche is recoverable as coordinated celltype co-abundance in HGCA scRNA-seq data, we examined compositional correlations among HGCA cell types mapped to HuBMAP’s definition of the niche, focusing on germinal center-associated states (**Supplementary Fig. 11**). Tolerogenic dendritic cells, follicle-associated resident macrophages, and reticular cell populations were positively correlated with this niche, confirming the coexistence of known follicle niche cell types alongside germinal center B cells. Thus, coordinated cell-type composition within the HGCA recapitulates the cellular architecture of a spatially defined intestinal niche despite loss of direct spatial information during tissue dissociation.

GC B dark-zone (DZ) and GC B light-zone (LZ) subtypes were both affected compositionally by radial tissue layer (DZ R2 =0.262, LZ R2=0.214). Since follicles form discrete structures within the intestinal wall, punctate biopsies and slices of surgical resections are expected to generate samples with either detectable or absent follicle signatures. We found that GSVA-based GC B gene-module scores (Methods) showed strong bimodal distributions in both biopsies and resections, particularly in the ileum by Gaussian mixture modeling and Ashman’s D (ileum ΔBIC=93.8, D=4.12; colon ΔBIC=37.4, D=4.04) (**Fig. 3i left**). Using combined GC B LZ and DZ abundance (GC B LZ + GC B DZ > 2 cells per sample) to estimate follicle capture, detectable follicle structures were present in 44% of ileal biopsies and 40% of colon biopsies, compared with 18% and 16% in ileal and colon resections (**Fig. 3i right**). A GC B abundance threshold scan placed detection rates closer to the literature’s expected higher density in ileum rather than colon^49^, but the 3 cell cutoff produced a boolean follicle call per sample that agreed best with germinal center gene module GSVA results (**Supplementary Fig. 12a-c**). Among covariates, follicle capture rate was significantly associated with sample collection method (resection vs biopsy) and increased with number of cells captured per sample, demonstrating how both anatomical sampling and sequencing depth influence the probability of recovering spatially restricted tissue structures (**Supplementary Fig. 12d-f**).

Having defined specific anatomical niches compositionally, we next asked whether cellular communication analysis suggested compartment or niche-specific wiring. Across gut segments, stromal and capillary endothelial states formed stable communication hubs, whereas the centrality of lymphatic and arteriolar endothelial populations shifted along the proximal-distal axis (**Supplementary Fig. 13a**). Endothelial wiring was dominated by the ACKR1 chemokine sink on pre-venule capillary (PVC) and venular endothelium, receiving diverse chemokines from lymphoid, stromal, and follicle-adjacent senders, with a related deep perivascular motif of perivascular resident macrophages engaging PAC/PVC/arteriolar endothelia via chemokine-ACKR1 and S100A9-CD36 axes (**Supplementary Fig. 13b**). Lymphatic and medullary sinus endothelium preferentially expressed CCL21, which binds with CCR7 on migratory DCs and naive/follicular T cells (**Supplementary Fig. 13c**). Communication involving follicle-associated resident macrophages was distinct from this broader perivascular program. While perivascular macrophages primarily contributed chemokine signals towards ACKR1-expressing endothelium via CXCL8, follicle-associated cell types showed predicted interactions with lymphoid populations within the follicular niche, including B cell activating factor (TNFSF13B) signaling to GC B and Memory B cells and LGALS3 interactions with CD4 Tfr (**Supplementary Fig. 13d-e**). LGALS3-associated communication was also upregulated from a number of enterocyte subtypes to CD4 Tfh in samples containing follicles (**Supplementary Fig. 13f**). Together, these analyses showed that coordinated cell-type compositions in the HGCA can infer the capture of spatially discrete intestinal niches and connect their cellular architecture to niche-specific communication programs, extending anatomical information from dissociated scRNA-seq beyond individual cell identities to higher-order tissue organization.

### Mapping Crohn’s disease onto the HGCA reveals cellular remodelling and treatment-response signatures

To demonstrate how the HGCA facilitates rapid annotation of new datasets and places disease signatures within a standardized cellular coordinate system, we mapped 987,466 cells from terminal ileum and colon samples of the TAURUS inflammatory bowel disease cohort^50^ to the HGCA using scArches^51^ (**Fig. 4a**). TAURUS includes adults with Crohn’s disease and ulcerative colitis (UC) sampled before biologic treatment (anti-TNF therapy), together with limited healthy controls (n=3). ScArches mapping accurately projected HGCA identities across all four major cell lineages^52^, allowing the disease cohort to be analyzed using the same cellular nomenclature as the healthy reference (**Fig. 4b**). To make this workflow accessible beyond computational specialists, deployment of the HGCA reference models through ArchMap is planned^53^, to provide a rapid point-and-click route from processed single-cell data to reference mapping and cohort-level analysis. ArchMap deployment is pending. We evaluated label transfer performance by manually matching TAURUS labels to their nearest HGCA labels, then quantifying which cells obtained labels of higher resolution, lower resolution, or were reassigned to varying degrees across our cell type hierarchy. The TAURUS annotation comprised 109 homeostatic and disease-associated states, whereas HGCA mapping assigned 84 healthy reference identities. Of these, 42 corresponded to matched TAURUS labels, while 42 represented new standardized healthy subtypes, including rare or transitional populations (**Supplementary Fig. 14a-b**). For instance, ABCA8pos WNT2Bpos FOShi fibroblasts were predominantly mapped to lamina propria fibroblasts, C3hi RSPO3pos fibroblasts to subepithelial fibroblasts, and a subset of non-ileal PLCG2hi undifferentiated enterocytes to mid crypt colonocytes. Most annotation changes represented differences in resolution or reassignment between closely related populations, whereas major reassignments were concentrated among epithelial progenitor states.

**Figure 4.**
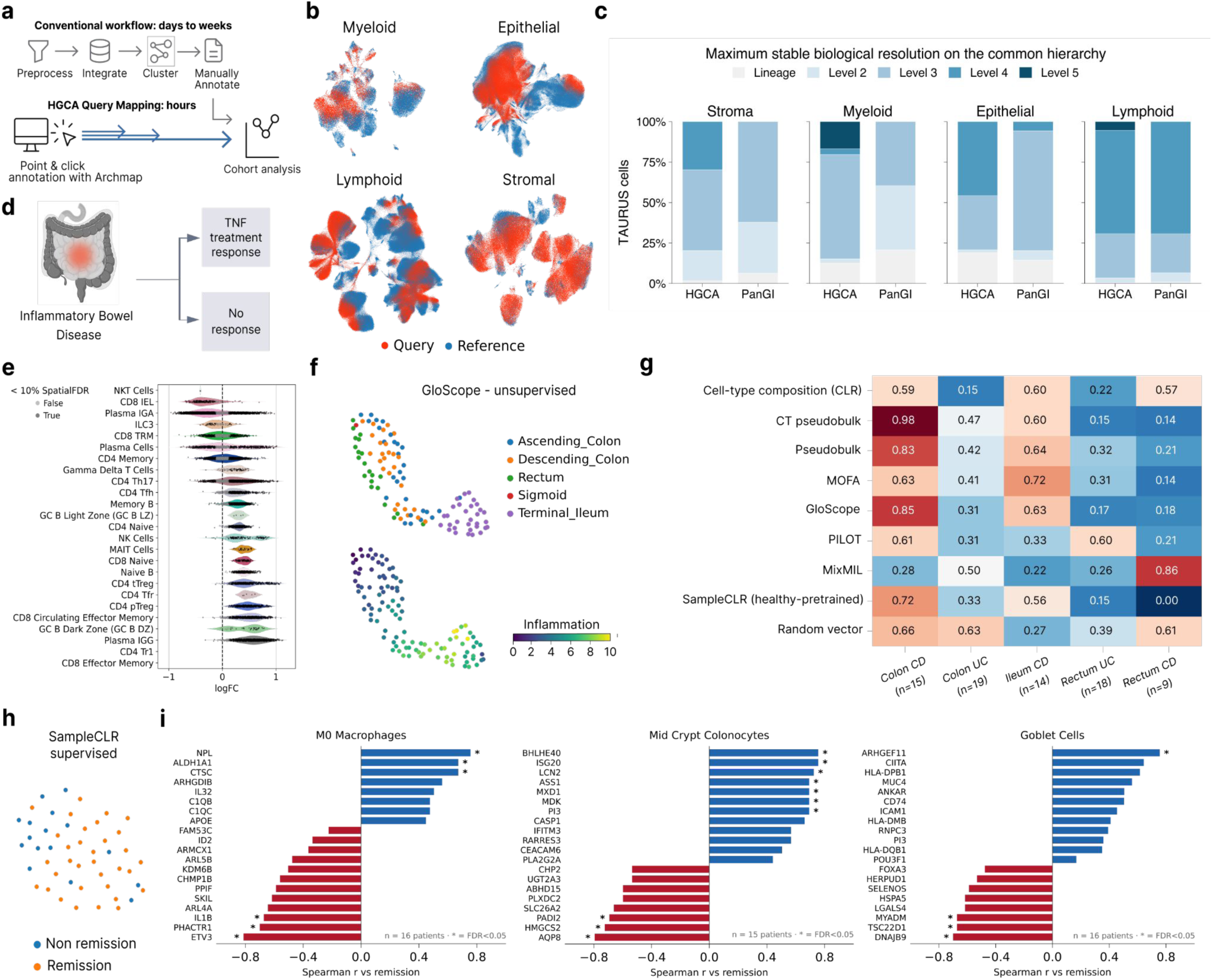
HGCA enables rapid modelling of IBD data. a) Comparison of the conventional single-cell analysis workflow with HGCA query mapping for rapid progression to cohort analysis. b) Lineage-specific UMAPs showing SCANVI mapping of TAURUS query cells onto HGCA reference cells. c) Maximum stable mapping resolution of HGCA labels to TAURUS. HGCA labels are higher resolution and more robust across constituent datasets. d) Schematic of the TAURUS inflammatory bowel disease cohort. e) Neighborhood-level differential abundance analysis of lymphoid cells of colon samples associated with inflammation. Distributions show neighborhood log fold changes grouped by transferred HGCA cell type, points individual neighborhoods. f) UMAP of unsupervised GloScope embedding of all samples from CD and control patients capturing anatomic location and inflammation. g) ROC-AUC for predicting eventual remission from pretreatment samples across sample embedding methods, tissue segments, and IBD subtypes h) UMAP of SampleCLR embedding supervised for remission status of pre-treatment samples, only i) Genes with highest correlation to remission status within cell types with high attention scores in the supervised sampleCLR model.

Recent benchmarking of single-cell foundation models has shown that learned cellular representations are dependent on the contexts of their constituent atlases and do not generalize well to new biological contexts, data modalities, and dataset composition^54^. To determine whether transferred identities are affected by the cellular, anatomical, and experimental contexts of the individual datasets of gut atlases, we rebuilt the HGCA and PanGI after dropping their shared healthy studies one at a time, remapped the same TAURUS cells, and measured the highest resolution cell type preserved across dataset hold-outs (“maximum stable biological resolution”; **Fig. 4c**). HGCA achieved a higher stable resolution across all cell type lineages, indicating that HGCA mapping is supported by more distributed evidence across its component datasets rather than being dependent on the particular context of one dataset. In cases where PanGI provides a lower resolution mapping, for example among absorptive epithelial cells, transferred identities using the HGCA were less stable, but higher resolution (**Supplementary Fig. 14c**). Importantly, this instability propagated into the disease analysis - after matching both references to the same biological resolution, TAURUS cell type compositions changed less under HGCA when reference studies were removed (**Supplementary Fig. 14d**).

To confirm the robustness and clinical relevance of downstream analysis results using the HGCA labels and coordinates, we performed compositional and cell state analysis of inflammation signatures and TNF-treatment response in intestinal tissue (**Fig. 4d**). In UC-affected colon and rectum, compositional analysis showed that Plasma cells shift from IgA-towards IgG-producing states (**Supplementary Fig. 15b**). Using Milo^55^ on the continuous inflammation score provided by the TAURUS study, we recovered inflammation-associated remodelling across multiple cellular compartments. GC Dark Zone B cells were enriched with increasing inflammation, consistent with expansion of lymphoid aggregates in intestinal inflammation^56^ (**Fig. 4e**). In the myeloid compartment, classical monocytes increased while homeostatic macrophages decreased with inflammation (**Supplementary Fig. 15c**).

Mapping new samples to the HGCA has the power to transform the HGCA from a descriptive atlas to a diagnostic tool. We examined whether HGCA-mapped data could support patient-level representations of disease and treatment-associated variation. Using patpy^57^, we compared sample-representation approaches based on cellular composition and gene expression profiles (**Supplementary Fig. 16a**). Expression-based representations captured biological variation associated with inflammation and anatomical location and supported prediction of remission following anti-TNF treatment, with particularly strong performance in colonic samples from patients with CD (**Fig. 4f-g**).

To identify cellular and molecular features underlying patient representations, we applied SampleCLR, a contrastive learning method for generating patient representations, supervised for remission status **(Fig. 4h**). SampleCLR assigns attention scores to each cell, allowing users to interpret the resulting representation with respect to a covariate of interest. We selected cell types based on their average attention score (**Supplementary Fig. 16b**), and filtered for genes with high correlation to per-cell attention scores as well as remission status within each cell type ( **Fig. 4i**). HGCA M0 macrophages were assigned one of the highest attention scores driving the embedding. SampleCLR identifies ETV3, PHACTR1, and IL1B being negatively, and anti-inflammatory NPL positively correlated with remission. High levels of C1QB and C1QC indicate that the identified cell population resembles the TAURUS study’s remission-associated macrophage cell state “C1Q^hi^ IL1^lo^ macro”. Within the epithelial compartment, mid crypt colonocytes with high levels of interferon signaling-related genes (ISG20, LCN2, PI3) and goblet cells with increased antigen presentation activity (CIITA, HLA-genes, CD74), confirm further remission-associated signatures suggested by the TAURUS study.

Together, these analyses demonstrate the rapid identification of disease-affected cell states and transcriptional signatures contextualized by their corresponding healthy cell types. The HGCA provides a reliable reference to organize disease-associated cellular remodelling, contextualize clinically relevant cohort-level variation, and connect patient-level phenotypes to reproducible cellular identities without requiring manual *de novo* annotation.

### HGCA mapping separates adult-state maturation from regional identity in organoids

Human intestinal organoids provide tractable experimental models for studying intestinal biology in health and disease, but their biological relevance depends on how faithfully they reproduce the cellular identities, regional specialization and maturation states of native tissue. Organoid fidelity is often assessed using broad cell classes or selected markers, potentially obscuring differences between maturation and regional identity. The Human Endoderm Organoid Cell Atlas (HEOCA)^58^ showed that organoid source and culture conditions influence intestinal cell composition and differentiation, establishing a framework for cross-protocol comparison. We reasoned that mapping these model systems to the higher resolution HGCA taxonomy could reveal which adult regional states are obtained, which remain immature or misplaced, and how organoid compositions approach native tissue with time in culture. We mapped 265,728 cells from 98 HEOCA gut organoid samples to the HGCA epithelial reference using SysVI^59^, retaining 192,017 cells with mapping confidence > 0.5 and 89 samples containing at least 500 confident cells (Methods; **Fig. 5a**). Mapping expanded 16 HEOCA labels into 28 HGCA identities, resolving stem and progenitor populations, crypt-villus absorptive states, tuft maturation, and regional BEST4 subtypes. Molecular condition and maturation time showed the strongest study-adjusted associations with composition, followed by publication, suggesting residual study-specific protocol or provenance effects not captured by available HEOCA metadata (**Fig. 5b**).

**Figure 5.**
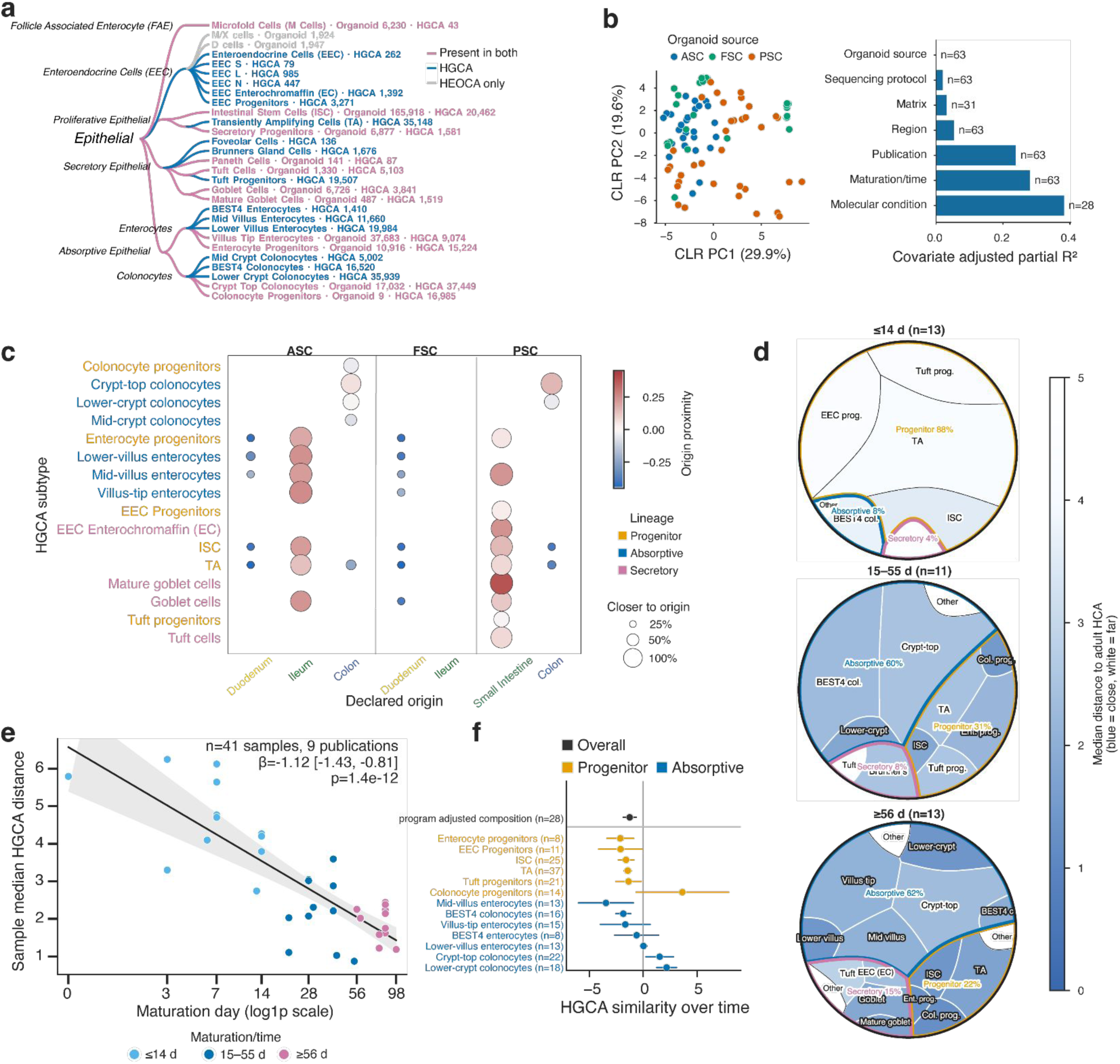
HGCA resolves cell subtype specific adult and regional correspondence in human gut organoids. a) SysVI mapping expands 16 HEOCA epithelial cell types to 28 HGCA identities among confidently mapped cells (mapping confidence > 0.5). b) PCA of CLR-transformed organoid compositions, colored by organoid source (left) and covariate-adjusted partial R2 from additive linear models. Bars show the unique compositional variance associated with each covariate; n denotes samples with non-missing metadata. ASC, adult stem cell-derived; FSC, fetal stem cell-derived; PSC, pluripotent stem cell-derived.. c) Segment of origin correspondence of HGCA epithelial subtypes by organoid source and declared segment of origin. Color denotes median origin-proximity score; point size denotes fraction of cells closest to declared segment of origin. d) Voronoi treemaps summarizing Small intestine PSC-derived organoid compositions at ≤14 days (n=13), 15–55 days (n=11), and ≥56 days (n=13). Tile area is proportional to mean cell-type CLR composition, and color denotes median distance to the nearest adult HGCA reference cell, with lower values indicating greater similarity. Colored outlines group progenitor, absorptive, and secretory identities. e) Median nearest-reference distance per PSC organoid plotted against log-transformed maturation day. The line and shaded band show fitted linear regression and 95% confidence interval. The reported coefficient is from a publication-adjusted linear model; p value from a two-sided t-test of the maturation-time coefficient. f) Per-cell type regression coefficients and confidence intervals, with the same model as **5e**. N denotes number of organoid samples contributing to each estimate.

To quantify regional fidelity, we examined whether mapped HEOCA cells were closer to adult HCGA reference cells from their declared segment of origin than to those from a different segment (**Fig. 5c**). Ileal identity was most consistently preserved, with 58.1% of adult stem cell-derived (ASC) and 57.1% of fetal stem cell-derived (FSC) ileal cells mapping closer to ileal references, compared with 32.6% of ASC colonic cells mapping preferentially to colon and 1.1% and 0.5% of ASC and FSC duodenal cells, respectively, mapping preferentially to duodenum. The limited duodenal specificity may partly reflect the smaller HGCA duodenal reference (8 donors; 13k cells). At cell-subtype resolution, 96.9-99% of ASC ileal villus enterocytes were ileum-proximal, with BEST4 enterocytes and several progenitor states also retaining ileal correspondence. Among PSC organoids, 66.6% of cells from small-intestine-targeted cultures were SI-proximal, whereas only 16.7% of cells from colon-targeted cultures were colon-proximal. Mature goblet, enterochromaffin, and mid-villus states showed the strongest SI correspondence, whereas crypt-top colonocytes were the most regionally faithful state in colonic PSC organoids. Differentiated absorptive and secretory states generally retained more segment-of-origin information than progenitors.

High-resolution region-specific cell type mapping also tested whether induction of a marker program reflected acquisition of the intended *in vivo* cell state. Childs *et. al*. showed that replacing EGF with EREG generates fetal enteroids containing BEST4+ enterocytes^60^. In the two HEOCA samples from this comparison, EREG increased HGCA BEST4 assignments from 29 of 3,318 confident cells (0.87%) to 152 of 3,184 (4.77%). However, 151 of 152 EREG-associated BEST4 cells mapped to BEST4+ colonocytes rather than BEST4+ enterocytes. These cells lacked expression of CFTR but were enriched for VSIG2, CA2, and HMGCS2, indicating induction of a BEST4-like program without acquisition of the expected duodenal identity. EREG was not recorded in the HEOCA molecular-condition metadata for these samples, illustrating how incomplete protocol annotation can conceal biologically meaningful culture effects (**Supplementary Fig. 17d**).

Longer PSC organoid protocols seek to carry pluripotent stem-cell derived intestinal tissue beyond its initial fetal-like state toward a more differentiated, adult-like intestinal epithelium^58^. Greater organoid-wide similarity to adult tissue can arise as differentiated lineages replace progenitors. We therefore tracked composition and nearest-reference distance together across SI-targeted PSC organoids (**Fig. 5d**). Early SI-targeted cultures (<14 days; n=13) were predominantly composed of progenitor states (84.7% progenitor, 7.9% absorptive, 3.7% secretory). Absorptive populations expanded at 15-55 days (n=11) and >55 days (n=13), accompanied by increasing representation of differentiated secretory and absorptive villus states (**Fig. 5d; Supplementary Fig. 17a**).

Although compositions point to a mix of colon and SI specific cell types in SI or colon-targeted PSC organoids with increased maturation time, across 41 PSC samples from nine publications, median nearest-reference distance decreased from 4.27 at <14 days to 2.22 at 15-55 days, and 2.00 at >55 days (Spearman ρ = -0.75; p=1.8×10^-8^; **Fig. 5e**). In a sensitivity analysis restricted to 31 samples from five publications containing multiple time points, the association persisted after adjustment for publication (β = -1.06/log(day+1); 95% CI [-1.37, -0.74]; p=4.3×10^-11^), with concordant effects within all five studies. Notably, increased adult fidelity was also evident within individual cell identities. Mid-villus enterocytes, enterocyte and EEC progenitors, ISCs, and TA cells each moved closer to adult HGCA references (**Fig. 5f**). EEC progenitors also acquired lineage markers, but only after early time points (**Supplementary Fig. 17d**). In contrast, colonocyte progenitors and crypt-associated colonocyte states moved farther from adult references, whereas BEST4+ colonocytes became more adult-like, indicating that prolonged culture did not uniformly resolve regional identity.Thus, HGCA mapping separates 2 dimensions of organoid fidelity that are often conflated, demonstrating that prolonged culture promotes acquisition of adult-like epithelial states while regional identity remains incompletely specified. Establishing both dimensions of fidelity will be important for disease modelling, where distinguishing disease-associated phenotypes from differences in epithelial maturation and regional or niche specification is essential for interpreting organoid-based perturbations and therapeutic responses.

## Discussion

The HGCA establishes a community curated reference framework for interpreting cellular variation from single-cell assays across the human gut. We engaged the community to generate i) a hierarchical consensus taxonomy of 94 intestinal cell types across five annotation levels, ii) harmonized anatomical, clinical, and technical metadata, and iii) a transferable reference mapping workflow that places gut datasets into the same framework. Together, these components provide a standardized coordinate system for interpreting intestinal single-cell datasets across studies, anatomical locations, and experimental platforms. This coordinate system characterized anatomical differences in cellular compositions, helping to guide future experimental design choices. We show that consensus identities generalize across healthy tissue samples, human intestinal organoids, and Crohn’s disease cohorts, enabling rapid and reproducible annotation, robust identification of disease-associated remodeling, and quantitative assessment of model system fidelity. The HGCA serves both as a reference of the healthy human gut and as a scalable analytical framework that enables new datasets, technologies, and biological knowledge to be continuously integrated into a shared community resource for studying intestinal biology in health and disease.

A defining feature of the HGCA is its transparent, community-governed annotation strategy. Using two international annotation jamborees and the Cell Annotation Platform (CAP), cell identity assignment became a transparent and versioned process that can be continuously refined as new biological knowledge emerges. Community consensus substantially increased the resolution and consistency of intestinal cell identities while improving transferability to independent datasets. Nevertheless, transfer performance remained lower for several cell populations, including neutrophils and exhausted CD8 T cells, indicating that future versions of HGCA will need to improve the representation of these cell states that are relatively rare in the HGCA. Future releases of the HGCA will therefore benefit from larger cohorts, improved sampling of transitional populations and continued refinement of cell identity definitions.

A second defining contribution of the HGCA is to anchor cellular identity within an anatomical coordinate system of the human gut. Cell composition and transcriptional programs varied reproducibly along both the longitudinal and radial axes of the intestine, demonstrating that anatomical location is a fundamental determinant of intestinal cell identity. Independent spatial transcriptomic data confirmed this variation by localizing consensus populations to their expected epithelial, stromal, vascular, and immune compartments. In the HGCA, differences in cellular composition reflected the depth and geometry of tissue sampling, with biopsies preferentially recovering superficial mucosa and full-thickness resections extending into deeper vascular, stromal, and outer-wall compartments. Although cross-study compositional differences are often obscured by study-specific sampling, processing, assay, and annotation choices,^61^ harmonized metadata further enabled reconstruction of higher order tissue organization, including lymphoid follicle capture, while segment-specific gene expression uncovered biologically meaningful distinctions within shared cell identities. Together, these findings establish anatomic location, sampling depth, and tissue provenance as essential coordinates for interpreting both the cellular compositions and molecular programs recovered by intestinal single-cell assays.

A major application of the HGCA is reference mapping analysis of new datasets. Analogous to the role of the human reference genome in sequence analysis, the HGCA provides a stable cellular and anatomical coordinate system that allows single-cell datasets to inherit accumulated knowledge of gut cell identity and anatomy at the point of mapping rather than requiring repeated *de novo* annotation. In the independent TAURUS IBD cohort, reference mapping supported immediate sample-level analysis of disease- and treatment-associated variation. Mapping human intestinal organoids similarly enabled simultaneous assessment of cell type representation, maturation, segment specification, and transcriptional proximity to primary tissue. Together, these analyses demonstrate that a standardized healthy reference provides a reproducible basis for comparing disease cohorts, perturbations, and experimental models while reducing the repeated annotation and harmonization that currently delay biological analysis.

The HGCA builds upon a growing collection of intestinal reference resources. HuBMAP established anatomically resolved spatial maps of the human intestine^8^, the Helmsley Gut Cell Atlas generated extensive clinically annotated IBD/CD datasets^10,13,24,26,29^, and PanGI demonstrated the feasibility of integrating single-cell datasets into a queryable reference atlas of the gastrointestinal tract^9^. Among these resources, PanGI is currently the only publicly available atlas designed for reference mapping of new datasets and therefore shares important conceptual goals with the HGCA. As a major contributor to the HGCA, PanGI also adopted the HGCA anatomical coordinate metadata to become more standardized as a response to reviewer comments, further promoting harmonization across community resources. However, PanGI integrates fewer datasets within individual intestinal segments, due to the cross-organ focus, and holds substantially lower stable cell type resolution when mapping new out-of-distribution (i.e. diseased) data. This distinction highlights that reproducible mapping and biologically informative resolution are separate properties of a reference, and that stable interpretation requires both. The HGCA extends these efforts through broader integration of healthy intestinal datasets, community-governed consensus annotation, harmonized anatomical and clinical metadata, and a unified reference framework that supports high-resolution annotation and robust transfer across independent healthy, disease and organoid datasets.

As the first official Human Cell Atlas reference for the human intestine, the HGCA is designed as a continuously evolving community resource. Its current release establishes a standardized anatomical and cellular framework for the healthy gut that can serve as a stable reference for future expansion. An important next step will be extending this framework across the full spectrum of disease, providing a baseline against which pathological remodeling can be measured and disease-specific modules that capture inflammatory, neoplastic, and regenerative states within a common cellular taxonomy, anatomical framework, and metadata standard. In this setting, diagnosis alone will be an incomplete description of biological context. Cellular phenotypes also reflect genotype, treatment history, comorbidities, exposures, and position along a disease trajectory^62–64^. Future HGCA disease atlases will therefore depend on richer, harmonized, privacy-preserving clinical metadata and sustained participation from clinicians and clinical data specialists. To support this transition, we defined 50 important protected clinical covariates prioritized for future collection (**Extended Data Table 4**), whose availability will be transparently tracked within HCA governance structures while protecting participant rights and confidentiality.

Realizing this vision will require overcoming the limitations of the current release. The HGCA v1 is dominated by adult mucosal tissue and underrepresents pediatric populations, upper gastrointestinal regions, and global population diversity, while reliance on dissociated scRNA-seq limits representation of neuronal, muscular, and other difficult-to-isolate populations. Continued expansion across populations, anatomical regions, and disease states, together with incorporation of spatial transcriptomics, single-nucleus, epigenomics, and proteomics, could place intestinal gut cell states within their native anatomical, molecular, and regulatory contexts. As future HGCA releases become increasingly multimodal, advanced computational approaches, including human gut foundation models, will be required to represent these complementary axes of biological variation jointly while preserving their cellular and anatomical context. The HGCA provides biological infrastructure for future studies of the human gut through a standardized cellular, anatomical, and metadata coordinate system, a diverse reference for rapid and robust mapping of new datasets^65^, evidence-linked annotations, validated models, and biologically grounded benchmark tasks that can support increasingly AI-enabled biological analysis^66^. Together, these advances enable reproducible interpretation of healthy tissue, disease variation, and experimental models while helping knowledge to accumulate around a common cellular nomenclature shared by the international gut research community.

## Supporting information

Supplemental Table 2: Theoretical cell types

Supplemental Table 3: Cell type covariate variance explained

Supplemental Table 4: HGCA Tier 2 metadata example

Supplemental Table 5: Germinal center GSVA

Author contributions

HGCA Gut Bionetwork contributors

## Acknowledgements

K.K, M.Z and M.J.A were supported by the Leona M. and Harry B. Helmsley Charitable Trust. J.O.M is a New York Stem Cell Foundation – Robertson Investigator. J.O.M was supported by the Leona M. and Harry B. Helmsley Charitable Trust, The Pew Charitable Trusts Biomedical Scholars, The Broad Next Generation Award, The Kenneth Rainin Foundation, The New York Stem Cell Foundation, and The Cell Discovery Network, a collaborative funded by The Manton Foundation and The Warren Alpert Foundation at Boston Children’s Hospital. D.K is funded by the Deutsche Forschungsgemeinschaft (DFG, German Research Foundation; Heisenberg grant, KO 4891/6-1, 496078873; KO 4891/5-1, 447562338), Helmholtz Initiative and Networking Fund of the Helmholtz Association (Helmholtz Young Investigator Group, VH NG 1716), The Leona M. and Harry B. Helmsley Charitable Trust, Else Kröner Fresenius Stiftung (Key Project 2019_A136), the Federal Ministry of Research, Technology and Space (Bundesministerium für Forschung, Technologie und Raumfahrt; BMFTR) as part of the German Center for Child and Adolescent Health (DZKJ) (funding code 01GL2406A), The Kenneth Rainin Foundation, and Care for Rare Foundation. M.D.L. was supported by the Chan Zuckerberg Initiative Foundation (grant CZIF2022-007488, Human Cell Atlas Data Ecosystem) and the Chan Zuckerberg Initiative DAF, an advised fund of Silicon Valley Community Foundation (grant 2021-235155). J.B. was supported by NIH K01DK140610 and the AGA-Bristol Myers Squibb Research Scholar Award in Inflammatory Bowel Disease (AGA2024-13-01).

This work has been made possible in part by Biohub (grant number CZIF2022-007488) through a grant from the Chan Zuckerberg Initiative Foundation.

C.L., B.W., V.A.S. are supported by the Helmholtz Association under the joint research school "Munich School for Data Science - MUDS". This research was funded in whole, or in part, by the Wellcome Trust [Grant number 220540/Z/20/A]. D.O.-S. was supported by grant 2023-221471 from the Chan Zuckerberg Initiative D.A.F. and by Wellcome grant 220540/Z/20/A (Wellcome Sanger Institute Quinquennial Review 2021-2026).

## Author contributions

Author contributions are listed in a Supplementary Information file.

## Ethics approval and consent

Intestinal biopsies for the generation of Xenium Spatial Transcriptomics were obtained upon written consent. The investigation was performed in accordance with current ethical and legal frameworks, and the study protocols were approved by the Institutional Review Boards at the LMU University hospital.

## Methods

### Dataset selection

The PanGI atlas^9^ healthy subset was subsetted to exclude gastric and esophageal datasets. Additional literature datasets were searched and prioritized by data availability, author response, and whether it filled a demographic gap. Datasets were further excluded if they performed poorly in the integration. These included one large-scale unpublished dataset, two single-nucleus datasets^8,17^, and several datasets using older or unique single-cell assays^6,14^. New fastq alignment was attempted on older datasets, after which they still performed poorly in integration. These datasets are prepared and ready for new attempts at integration in later versions.

### Metadata ingestion

Metadata requirements were defined in a central Google Sheet containing separate tabs for dataset-, donor-, sample-, and cell type-level fields, together with descriptions, examples, and permitted values. MetaManager used this Google Sheet along with Google Drive APIs to generate standardized metadata entry sheets directly from these definitions, including dropdown menus for controlled fields. Existing metadata was prefilled where publicly available before sheets were distributed to study contributors, allowing authors to focus on missing or ambiguous fields.

Completed sheets were retrieved programmatically and assessed for completeness and correctness using allowed-value lists and regular-expression rules for structured fields such as ontology identifiers. Validation results were summarized by dataset and metadata field as heatmaps and published to a shared web page, providing contributors and atlas coordinators with a continuously updated view of unresolved gaps and invalid entries. Maintaining the entry sheets, validation rules, and project-wide tracking display in a shared system enabled rapid feedback, transparent attribution of outstanding metadata, and consistent assessment of readiness across successive atlas object versions.

### Dataset harmonization

Dataset, donor, and sample level metadata were harmonized before integration and datasets were individually uploaded without filtering to the HCA portal. The fields used in the present analysis included dataset, donor, and sample identifiers, anatomical gut segment, sampled site condition (disease adjacent versus healthy), radial tissue layer, collection and preservation methods, assay, sequenced fragment, gene annotation version, sex, and decade age ranges. Harmonized values and unresolved entries were tracked through the metadata tracker; unknown values were retained for summaries but excluded from contrasts that required a defined category. Contrasts were marked as having support only when at least one dataset contained two categories represented by at least five samples each.

### Quality Control

Quality control (QC) was performed following a two stage strategy. Before integration, a permissive filtering using lineage specific QC metrics was applied to allow potential detection of cell populations with lower QC metrics such as granulocytes (myeloid (n_counts > 300, n_genes > 120, MT_fraction < 0,25), epithelial (n_counts > 300, n_genes > 200, MT_fraction < 0,6), stromal (n_counts > 300, n_genes > 200, MT_fraction < 0.25), lymphoid (n_counts > 300, n_genes > 200, MT_fraction < 0,25)). After integration, cell clusters were evaluated carefully for being doublets or low quality cells and excluded for downstream analysis.

### Integration and clustering

Candidate integration models and batch keys were compared using scIB metrics for biological conservation and batch correction implemented in scAtlasTb^38^. scVI (ver 1.3.0) with donor identifier as batch key was selected for the final integration (non-default parameters: n_latent=30, n_hidden=128, n_layers=2, dispersion=’gene-batch’, dropout_rate=0.15). All-cell integration was performed for the purpose of visualization with UMAP. Epithelial, lymphoid, myeloid, and stromal lineages were iteratively clustered for cell type annotation. Leiden clusters were inspected at two resolutions, and marker genes were calculated across lineage clusters and within each cluster’s sub-clusters to provide information to expert annotators.

### Software developed

Project software includes: the metadata library used to validate contributor metadata and expose missing fields. Analysis code and data tables for each figure are included with the manuscript. Release identifiers and repository URLs are included in the Code Availability section.

### Cell type hierarchy

The R software ARBOL was used (Code Availability) for tree representations of cell types throughout the text. HGCA identities were represented in a five level hierarchy spanning broad lineage to high resolution cell subtype. Candidate labels were linked to canonical and assay specific markers, literature evidence, and Cell Ontology terms. Integrated lineage objects were uploaded to the Cell Annotation Platform (CAP), expert responses were reviewed, and accepted changes were incorporated into the v1 taxonomy while retaining the pre-CAP v0 labels. CAP responses were collected using REST API in python.

#### Hierarchical comparison of original and consensus annotations

Original author labels were manually mapped to their closest node in the five-level HGCA taxonomy before comparison with v1 annotations. Each label was represented by its path from the lineage root to its terminal taxonomy node. Comparisons were classified as:

**Refined**: when HGCA v1 label was a strict descendant of author label
**Unchanged:** when both labels terminated at the same node
**Coarsened:** when HGCA v1 label was an ancestor of author label
**Changed branch:** when the two paths diverged below their last shared ancestor

Cell, sample, donor, and dataset support for each v1 identity was computed from the final, filtered integrated objects. Heatmaps in **Fig. 2** additionally summarize presence in source author annotations and PanGI, CAP response counts, dataset prevalence, and compositional enrichment by gut segment, radial layer, and collection method (resection vs biopsy).

### Differential Expression Analysis

Differential expression (DE) was performed with samples pseudobulked with scanpy (>10 cells/sample) per hgca_celltype_v1 celltype label unless indicated otherwise. pyDESeq2 was used with default parameters and

formula = ∼ dataset_id + covariate

Where dataset_id is included as a blocking factor to control for dataset-specific expression differences.

#### Power analysis

Log_2_ fold changes and standard errors from pseudobulked DESeq2, as described above, contrasting ileum vs colon samples, were filtered for genes with Benjamini-Hochberg-adjusted p < 0.05 and absolute log_2_ fold change > 0.5 from the all-cells object. For each gene passing thresholds, standard errors were scaled to sample size. Gene-level two-sided Wald power (α=0.05) was computed from scaled standard errors and estimated log_2_ fold changes. Mean detection probability per gene (+/- s.d.; mean Wald power), was calculated at balanced sample numbers per segment from 8 to 120 samples, including the published-only and published-plus-contributed sample sizes for each cell type.

### Compositional Analysis

Per sample cell type counts were calculated within each lineage, given a 1 pseudocount and transformed using the centered log ratio (CLR) transformation:

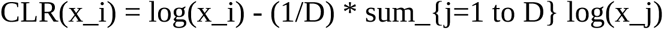

Where x_i_ is the pseudocount adjusted count of cell type i and D is the total number of cell types. Composition was calculated independently within each lineage unless otherwise noted.

### Compositional correlations

Sample-level cell type compositions with CLR were correlated across all samples in the HGCA with Spearman rank correlations. Cell types were included if detected (>2 cells) in >11 samples. P-values among tested pairs were adjusted with Benjamini-Hochberg false discovery rate (FDR). Types were ordered first with a curated Inner Follicle member block, followed by hierarchical clustering of the remaining types.

### Sample embeddings

Single-cell sample embeddings were calculated with multiple methods, in multiple settings with HGCA labels.

#### Crohn’s sample embeddings

Patpy^58^ was used for Taurus CD embeddings. For Taurus analysis, seven representation methods were compared: centered log-ratio (CLR) transformed compositions, sample and celltype pseudobulk, MOFA, GloScope, PILOT, MixMIL, against a random baseline.

#### Organoid sample embeddings

CLR compositional PCA was used to compare organoid samples with HGCA labels to uncover important axes of variation contributed by harmonized metadata.

### Covariates Analysis

CLR transformed cell counts were summarized using ordinary least squares (OLS) models and incremental partial R^2^/ω^2^. Omega squared was defined as:

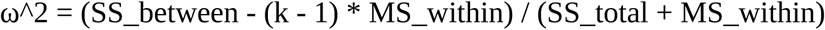

where k is the number of represented categories. Negative estimates were bounded at zero and estimates greater than 1 were bounded at 1. Unknown or unreported metadata values were excluded from the analysis of that covariate. Biopsy versus surgical resection (sample_collection_method) and full thickness EPI_LP_MUSC (radial_layer) versus all other radial annotations were tested with two-sided Mann-Whitney U tests, requiring at least five samples per group with FDR correction of p-values. Expression analyses used sample level pseudobulk log2(Counts per million). Depth associated differential expression was estimated with log2(CPM) ∼ group.

For each cell type and categorical covariate, one-way ω^2^ was calculated from between-category and residual sums of squares and bounded to the interval 0-1. Kruskal-Wallis tests were used to compare estimates. Pairwise category contrasts used two-sided Mann-Whitney U tests when each category contained at least five samples. Age range analyses used the ordered range midpoint and sample level Spearman correlation. FDR adjustment was applied across cell types within each segment, covariate, and contrast family.

Matched composition and expression effect sizes were calculated per sample for each cell type and aggregated to the lineage level using sample-count weights. The biological:technical statistic was defined as the ratio of the mean lineage ωZ for the gut segment to the mean lineage ωZ for the dataset identifier. Radial sampling depth was evaluated using two contrasts: biopsy versus surgical resection, and full thickness samples annotated EPI_LP_MUSC versus other radial annotations, excluding mesentery and appendix. Ileum and colon met the five sample thresholds for statistical tests, proximal segment contrasts are displayed as averages. BEST4 segment marker summaries report the percentage of cells expressing each gene and mean log1p expression normalized to 10,000 counts per cell.

#### Organoids covariates analysis

Metadata associations with organoid composition were quantified using multivariate ordinary least squares (OLS) models of compositional principal components. For each metadata variable, a full model containing that variable plus an adjustment set was compared with a reduced model omitting the variable of interest. Adjustment sets were as follows:

- Publication: organoid source, region, maturation/time
- Organoid source: region, maturation/time
- Region: organoid source, maturation/time
- Maturation/time: organoid source, region
- Matrix: organoid source, region, maturation/time
- Molecular condition: organoid source, region, maturation/time, matrix
- Sequencing protocol: organoid source, region, maturation/time

Covariate-adjusted partial R^2^ was calculated from the reduction in residual sums of squares across all components:

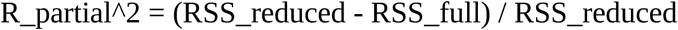

Models were fitted using samples with complete observations for all variables included in that comparison. Consequently, sample size varied across metadata variables and is reported in the figure (n=63 for source, region, time, publication and sequencing protocol; n=31 for matrix, n=28 for molecular condition).

### Germinal center (GC) lymphoid module detection

Analyses were restricted to ileum and colon samples from healthy or adjacent tissue. A GC-associated lymphoid module was defined from CAP taxonomy marker genes for germinal-center B cells (light and dark zone), expanded with the top Wilcoxon marker genes for these cell types within the lymphoid lineage object. Sample-level expression was obtained by summing raw UMI counts across all lymphoid cells within each sample_id (minimum 20 lymphoid cells), converting to log_2_(CPM + 1), and scoring the GC-module gene set with Gene Set Variation Analysis as implemented in decoupler (run_gsva; Gaussian kernel; minimum five genes per set). These genes are a GC B program built from CAP taxonomy markers for germinal-center / dark-zone / light-zone B cells (plus curated GC staples such as AICDA, BCL6, RGS13, CXCR4, CD83, MME, LMO2, SUGCT), expanded with top Wilcoxon marker genes from atlas GC B cells, spanning B-cell identity, MHC-II presentation, proliferation, and zone-associated GC programs (**Extended Data Table 5**).

#### Bimodality of GC capture

To test whether module enrichment was consistent with discrete presence versus absence across libraries, sample GSVA scores were fit to one- and two-component Gaussian mixture models (scikit-learn; five random initializations k=2). Difference in Bayesian information criteria (BIC) > 10 between these two models were used to identify strong evidence for bimodality as:

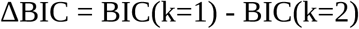

Ashman’s D was used to corroborate Bayesian statistics, where D > 2 suggests clear separation of normal distributions, with

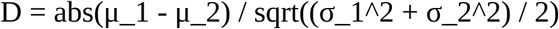

Where μ_1, μ_2 are the means and σ_1_,σ_2_ are the standard deviations. Distributions with both ΔBIC > 10 and D > 2 were interpreted as bimodal. Statistics were computed within ileum samples and within colon samples, within biopsy samples, and within resection samples.

### Composition based GC detection

Independently of GSVA, a sample was called GC B positive if it contained at least 3 cells annotated as GC B LZ or GC B DZ. Detection rates were stratified by gut segment (ileum, colon) and collection method (endoscopic biopsy versus surgical resection). Wilson 95% confidence intervals were calculated for binomial proportions. Biopsy versus resection rates within each segment were compared by two-sided Fisher’s exact tests; P values were adjusted for two tissues within each cutoff with Benjamini Hochberg FDR.

#### Threshold scan and GSVA concordance

Detection rates were computed for k = 1…40, stratified by segment and pooled, with Wilson score confidence intervals. Sample-level GSVA scores for Germinal Center (GC) modules and related follicle programs were taken from lymphoid lineage pseudobulk GSVA (explained above in GC module detection). For each k we summarized (i) the mean GC-module GSVA difference between follicle+ and follicle-samples and (ii) concordance of the composition call with GC module GSVA > 0 (Youden index = sensitivity + specificity - 1). The cutoff maximizing this concordance in the window k in [3, 40] was k = 3 so all main-figure analyses use k=3

## Covariate variance and confounding

For each metadata covariate we fit a univariate logistic model of the follicle call and reported McFadden’s pseudo-R^2^. Categorical predictors used the saturated within-level model (exact Bernoulli likelihood), continuous log1p(total cells/sample) used logistic regression. Contribution after study was the increase in pseudo-R^2^ when the covariate was added to an additive model that first included study as a covariate (no study x covariate interactions). Pairwise confounding among categorical covariates was summarized with Cramer’s V on contingency tables (levels with n < 5 excluded).

### Mixed-effects logistic model

We modeled follicle call as glmer(gc ∼ sampled_site_condition + collection_method + tissue_level_1 + scale(log_total_cells) + (1 | dataset_id) + (1 | donor_id), binomial family, Laplace approximation, bobyqa optimizer) Fixed-effect contrasts were disease-adjacent versus healthy, resection versus biopsy, and colon versus ileum. Odds ratios and Wald 95% CIs are reported.

### Identification of disease signatures

#### Enriched cell states in inflammation

Following benchmark results on how to best identify cell populations of interest in mapped scRNA-seq data, Milo^55^ was run on the shared scANVI^72^ embedding subsetted for the TAURUS data only. As the variable for enrichment testing the inflammation score per sample provided by the original TAURUS data was used. The mapped HGCA cell type label was associated with inflammation enriched neighborhoods.

#### Pretreatment anti-TNF response prediction

Samples were restricted to pretreatment biopsies with binary clinical outcome labels (remission vs non-remission after anti-TNF) and evaluated each tissue x disease context separately (ileum CD, colon CD, rectum CD, rectum UC, rectum UC). Predictive models were built on sample embeddings as described above. Outcome probabilities were obtained from distance-weighted k-nearest-neighbor classification on the embedding (k=5). A nested leave-one-patient-out cross-validation design was used for Area under the Receiver-operating characteristic Curve (ROC-AUC): for each patient held out, highly variable gene selection, scaling, PCA, representation fitting, and neighbor selection were performed using only the remaining patients in that context. All pretreatment biopsies from the held-out patient were excluded together, and multiple biopsies from one patient were aggregated to a single patient-level feature vector before prediction. For CT-pseudobulk, PCA scores were computed per cell type present in at least 3 training patients and at least 50 cells, and concatenated across cell types. Prediction (i.e. discrimination) was summarized by ROC-AUC.

### Label transfer

#### Leave-one-dataset-out (LODO) Benchmarks

To evaluate whether CAP revisions improved HGCA annotations, we performed a leave-one-dataset-out (LODO) cross validation benchmark using lineage-specific SCANVI models with sample ID as batch key. The benchmark was run independently for each major lineage: epithelial, lymphoid, myeloid, and stromal. For each held out dataset, all cells from that dataset were excluded from SCANVI model training, and the remaining datasets were used to train a reference model. The held out dataset was then mapped back into the reference latent space with labels masked, allowing predicted labels to be compared to the atlas integrated annotations. This procedure was repeated using pre (v0) and post (v1) CAP revision high resolution cell type annotations. The healthy subset of the PanGI atlas^9^ was divided into the same four major lineages: epithelial, lymphoid, myeloid, and stromal. LODO cross validation was performed independently within each lineage. In each fold, all cells from the held out dataset were excluded before fitting a separate SCANVI reference model to the remaining PanGI data. The held out cells were then mapped into the reference latent space with labels masked, and predicted labels were evaluated against the PanGI reference annotations using the F1 aggregation described below. Prediction performance was quantified using F1 score for each held-out dataset and label set. F1 scores were summarized by averaging F1 per cell type to give equal weight to rare and abundant cell types, then averaging cell type F1’s per label set / dataset to arrive at one F1 score per held out dataset. The v0, v1, and PanGI benchmarks used the same lineage-specific held out dataset design. Performance was calculated per cell type within each held out dataset and summarized as macro F1 by averaging F1 across rare and abundant cell identities to give them equal weight.

#### Organoid label transfer

Cell type label transfer was performed using scVI and sysVI^59^ with k-nearest neighbors (kNN) classification (k=30, distance weighted) using the Epithelial lineage object as reference. The reference dataset was used to train an scVI model with sample_id specified as batch covariate. The query dataset was mapped onto the reference latent space using scArches, and the resulting query model was further trained for 50 epochs. For SysVI integration, reference and query datasets were concatenated, and HVGs were selected based on the intersection across datasets. The sysVI model was trained on the concatenated data with sample_id as batch. For each query cell, predicted labels, confidence scores (defined as maximum output label probability across each cell’s probability distribution across cell types), and Shannon entropy values (of the label mapping probability distribution) were computed. Cells with confidence < 0.5 were assigned unknown and excluded from analysis requiring transferred identities.

#### Visium data label transfer

To transfer single-cell transcriptomic profiles to Visium spatial transcriptomics data, we performed spatial mapping using cell2location (v0.1.4)^67^. For each cell type, we generated a scRNA-seq reference from the HGCA scRNAseq dataset by down-sampling to at least 5,000 cells per cell type. Prior to running cell2location, we filtered genes using the following thresholds: cell_count_cutoff = 5, cell_percentage_cutoff2 = 0.03, and nonz_mean_cutoff = 1.12. In the main cell2location model, we explicitly accounted for technical variation by modeling batch effects using sample_id. In addition, we included categorical covariates corresponding to donor_id and assay to further remove batch and other technical confounders. Model training and subsequent inference produced spatial cell-type abundance estimates for the Visium samples.

For downstream analyses, we inferred cellular compartments using factor analysis implemented in cell2location via the feature co-localized cell-combination model, which applies non-negative matrix factorization (NMF). We evaluated the number of factors over a range of 10–20. The resulting factor loadings were then used to represent the relative contribution of each inferred compartment and to delineate tissue zones/niches in the spatial context.

#### Xenium datasets

Two Xenium datasets from the Bionetwork were considered in this study using the 10X colon gene panel and the Prime 5K Human Pan Tissue & Pathways Panel. Xenium transcripts were assigned to cells with Segger (v0.1.0, dpeerlab/segger) and using cells_min_counts=5 for the 372-gene panel and otherwise default parameters. Both panels (372-gene colon panel and 5,000-gene custom panel) were processed identically. Each sample was segmented with a gene-gene correlation prior (--gene-corr-reference-path) derived from matching dissociated single cell data (including genes ≥ 100 counts, gene_missing_strategy=remove).

Per-sample Segger outputs were aggregated into a single AnnData (outer join on genes). Following Xenium best practice^49^. A hard floor of ≥ 10 counts per cell, a gene filter (≥ cells per gene), and removal of control/blank probes were used for cell filtering. Counts were library-size normalized to target_sum = 100 and log1p-transformed; raw counts were retained.

Denoising was done using resolVI^68^ (scvi.external.RESOLVI), an scvi-tools model for imaging-based spatial data that jointly corrects segmentation spillover and batch effects. It was trained on raw counts (with batch_key being processing batch, n_latent=10, max_epochs=50 for the 372-gene panel and 100 for the 5,000-gene panel, no early stopping, train_size=1.0). The spillover-corrected expression layer is used for visualizations. The background in presented images is the combination of the four channels captured by the cell segmentation kit.

#### Disease data label transfer

For disease label transfer, scANVI mapping was performed using fully trained scANVI models per lineage with the same parameters as the benchmark. See codebook.

#### Atlas Mapping Stability

Lineage-specific scANVI models were retrained for HGCA and PanGI after omitting one study at a time from each reference. Omits were restricted to studies shared by name between atlases per lineage:

- Stroma: Kinchen2018, Lee2020, Martin2019, Uzzan2022, Yu2021
- Myeloid: Dominguez2022, Huang2019, Lee2020, Martin2019, Uzzan2022, Yu2021
- Epithelial: Huang2019, Lee2020, Martin2019, Parikh2019, Uzzan2022, Wang2020, Yu2021
- Lymphoid: Dominguez2022, Huang2019, Jaeger2021, Kinchen2018, Lee2020, Martin2019, Parikh2019, Uzzan2022, Wang2020, Yu2021

The same TAURUS lineage cells were mapped to each model to obtain label probabilities and high-resolution label predictions. Models used 4,000 HVGs; n_latent = 30; n_layers = 2; negative-binomial likelihood; 10 training epochs; batch size 256; no early stopping. Stromal analysis used 10 full reference seeds and three seeds for each of five shared-study omissions; myeloid, epithelial, and lymphoid analyses used three full-reference seeds and one seed for each shared-study omission.

#### Maximum Stable Biological Resolution

Mapping label predictions were placed on a common hierarchy defined by the HGCA v1 taxonomy. PanGI labels were mapped to their closest HGCA label using the same HGCA-PanGI matching table used to define PanGI presence for each HGCA label in **Fig. 2c**. For each query cell and hierarchy node v, support was defined as the fraction of mappings in which the predicted label was v or one of its descendants. The maximum stable biological resolution of each cell was defined as the most specific node supported across ≥90% of mappings (effectively requiring exact label matches across all left-out datasets, to be supported since only at most 10 datasets were left out; in stroma, since 3 replicates were performed for each of the 5 datasets, 14/15 had to agree, for lymphoid, 9/10 had to agree). Information specificity was a continuous measure of the resolution at each node:

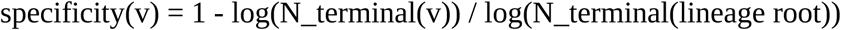

Where N_terminal_(v) is the number of terminal HGCA leaves descending from node v within that lineage. Terminal identities therefore have a specificity of 1, whereas broader identities containing multiple terminal cell types have progressively lower specificity. For visualization, maximum stable nodes per cell were grouped as lineage, level 2, level 3, or highest resolution.

### Organoid source nomenclature

Organoid samples in HEOCA are annotated by stem-cell origin as adult stem cell-derived (ASC), fetal stem cell-derived (FSC), or pluripotent stem cell-derived (coded “IPS” in the HEOCA metadata table). We refer to the latter as PSC-derived throughout, as it contains both human embryonic stem cell (h9 line) and induced pluripotent stem cell (iPSC)-derived organoids.

### Organoid *in vivo* proximity analysis

For each PSC derived organoid cell, nearest-reference distance was calculated in the SysVI latent space as the Euclidean distance to the single nearest healthy HGCA reference cell. Sample-level *in vivo* distance was defined as the median of distances across all query cells in the sample. Lower values indicate greater correspondence to adult HGCA epithelial transcriptomes.

Anatomical correspondence was quantified for query cells with mapping confidence of at least 0.5 that also passed a within-identity nearest reference distance outlier filter (z-score < 3.5). For each organoid cell, the distance to HGCA reference cells from the reported segment of origin, d_origin, and the distance to the nearest alternative segment d_alt were calculated as the mean Euclidean distance in the SysVI latent space (X_sysVI) to up to 50 nearest healthy HGCA reference cells from that tissue.

Relative segment of origin proximity was calculated per cell as

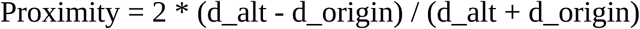

so that positive values indicate greater proximity to the reported segment of origin, zero indicates equal proximity, and negative values indicate greater proximity to another segment. For visualization, sample-celltype combinations with fewer than 50 non outlier cells were excluded. Median proximities and fraction of cells closer to the reported segment were then summarized across sample-celltype combinations.

### Organoid maturation analysis

Transferred HGCA epithelial cell types were grouped into three branches based on lineage descent and maturity: progenitor, differentiated absorptive, and differentiated secretory groups. BEST4 enterocytes and BEST4 colonocytes were classified as differentiated absorptive states. For each PSC-derived organoid sample, group fractions were calculated among confidently mapped epithelial cells. Associations of raw fractions with maturation time were evaluated using OLS regression HC3 with standard errors used for 95% confidence intervals. Numeric reported culture times were represented as days or log(1+day), and publication was included in the regression model as a categorical covariate.

Maturation models included 41 PSC-derived organoid samples from 9 publications with reported numeric culture times. The cross-publication association was estimated with an OLS regression model for aggregated sample-level median distances described in *proximity analysis*. Publication-adjusted analysis was restricted to 31 samples from 5 publications containing more than one maturation timepoint by adding publication category to the model. Regression coefficients, two-sided P values and 95% confidence intervals were calculated using HC3 standard errors. All five eligible publications showed negative estimated slopes.

### Gene ontology

Cell segment origins were classified as ‘small intestine’ if the cell originated from duodenum, jejunum, or ileum. Cells were classified as ‘large intestine’ if they originated from the cecum, ascending colon, transverse colon, descending colon, sigmoid colon, or rectum. Cells from the appendix and the mesentery were excluded. For each segment (i.e., small intestine, or large intestine), pseudo bulk analysis was performed for each cell type on a per donor level. Cell types with less than 3 unique donors per segment were excluded from the analysis. Given that colonocytes are the specialized enterocytes located in the large intestine, cells that were labeled as Colonocytes or Enterocytes were relabeled as Absorptive Epithelial to enable differential analysis between the enterocytes of the small vs large intestine. DESeq2 (v.1.42.1) was used to perform differential analysis on the pseudo bulk data for each cell type using a design formula ∼segment. gProfiler2 (v.0.2.3) was then used to perform enrichment analysis on the significant genes (adjusted p-value <0.05) using Gene Ontology Biological Process (GO:BP), on the hsapiens organism using all annotated genes as the background. GO terms were filtered to remove term sizes <5 and >350. GO terms were manually curated to identify the broad categories i.e., Immune Response, Metabolism, Mitochondrial Respiration, Digestion and Absorption, Transport And Secretion, Cell Cycle, Stress Response, Adhesion And ECM, Glycosylation, and Translation. Terms that were not assigned to these categories were classified under a category of “Other”. For visualization, GO terms were filtered to include the top significant term (based on p-value) per cell type and per direction.

### Cell-cell Communication Inference

Ligand–receptor (L–R) communication was inferred with LIANA rank aggregate. Ligand– receptor communication was inferred separately in each major gut segment (tissue_level_1: duodenum, jejunum, ileum, colon) using LIANA (v1.7.1) in Python (scanpy v1.11.5, anndata v0.11.4, pandas v2.3.3, numpy v2.2.6, scipy v1.14.1; Python 3.11) on the HGCA all-lineages object. Analyses were restricted to cells annotated as disease = normal. Mesentery and appendix were excluded. The low-quality catch-all state “Epithelial” was removed from downstream tables.

Counts were normalized to 104 and log1p-transformed. Genes detected in fewer than five cells were discarded. To keep runtimes tractable while preserving celltype representation, each segment was cell type stratified subsampled to at most 12,000 cells (segments with fewer than 2,000 cells were skipped). LIANA’s default filters were used as well.

Communication was scored with LIANA’s rank_aggregate consensus over CellPhoneDB, Connectome, log2FC, NATMI, and SingleCellSignalR using OmniPath consensus L-R (resource_name = "consensus"). Interactions were retained only if both ligand and receptor were expressed in at least 10% of cells in the sender and receiver populations, respectively (expr_prop = 0.1). Permutation testing used 1,000 permutations (n_perms = 1000). Method ranks were aggregated by Robust Rank Aggregation (aggregate_method = "rra"), yielding a magnitude consensus rank (magnitude_rank) and a specificity consensus rank (specificity_rank).

#### Cross-segment ensemble robust interactions

To identify interactions supported across the gut axis, we multiplied magnitude rank and specificity rank and log_10_ transformed this quantity. An interaction was defined as ensemble-robust if it was top-ranked (top 1% magnitude rank) in all four segments.

## Data availability

The spatial transcriptomics data are publicly available on Zenodo at https://doi.org/10.5281/zenodo.22148916.

## Code availability

Metadata manager: https://github.com/CellDiscoveryNetwork/MetaManager

Tutorial code book: https://github.com/HCA-integration/hca-gut-atlas-tutorial (tutorial updates pending).

Additional downstream analysis repo: https://github.com/HCA-integration/hca-gut-atlas-downstream

ARBOL: https://github.com/jo-m-lab/ARBOL

Patpy: https://github.com/lueckenlab/patpy

## Gut Cell Atlas Crohn’s Disease Consortium

Carl A. Anderson^1^, Mark J. Arends^2^, Anindita Basu^3^, Guy E. Boeckxstaens^4^, Eugene C. Butcher^5^, Lori A. Coburn^6,16^, Shahida Din^2,15^, Jellert T. Gaublomme^7^, Kathryn E. Hamilton^8^, Narayanan Kasthuri^3^, Bartha M. Knoppers^9^, Dermot P. B. McGovern^10^, Jeffrey R. Moffitt^11^, Andreas E. Moor^12^, Peng Qiu^13^, John F. Randell^14^, Keith T. Wilson^6^

1. Wellcome Sanger Institute, Wellcome Genome Campus, Hinxton, Cambridge, UK.
2. Institute of Genetics & Cancer, University of Edinburgh, Edinburgh, UK.
3. University of Chicago, Chicago, IL, USA.
4. KU Leuven, Leuven, Belgium.
5. Palo Alto Veterans Institute for Research, Palo Alto, CA, USA.
6. Vanderbilt University Medical Center, Nashville, TN, USA.
7. Columbia University, New York, NY, USA.
8. Division of Gastroenterology, Hepatology, and Nutrition, Department of Pediatrics, University of Pennsylvania Perelman School of Medicine and Children’s Hospital of Philadelphia, Philadelphia, PA, USA.
9. McGill University, Montreal, Quebec, Canada.
10. Cedars-Sinai Medical Center, Los Angeles, CA, USA.
11. Boston Children’s Hospital, Boston, MA, USA.
12. ETH Zürich, Basel, Switzerland.
13. Georgia Institute of Technology, Atlanta, GA, USA.
14. Broad Institute of MIT and Harvard, Cambridge, MA, USA.
15. Edinburgh IBD Unit, Western General Hospital, Edinburgh, UK.
16. Veterans Affairs Tennessee Valley Healthcare System, Nashville, TN, USA.

