## Supplementary Figures 1-17 for "The Human Gut Cell Atlas v1.0: A Unified, Community-Built Reference of Cellular Diversity Across the Gut"

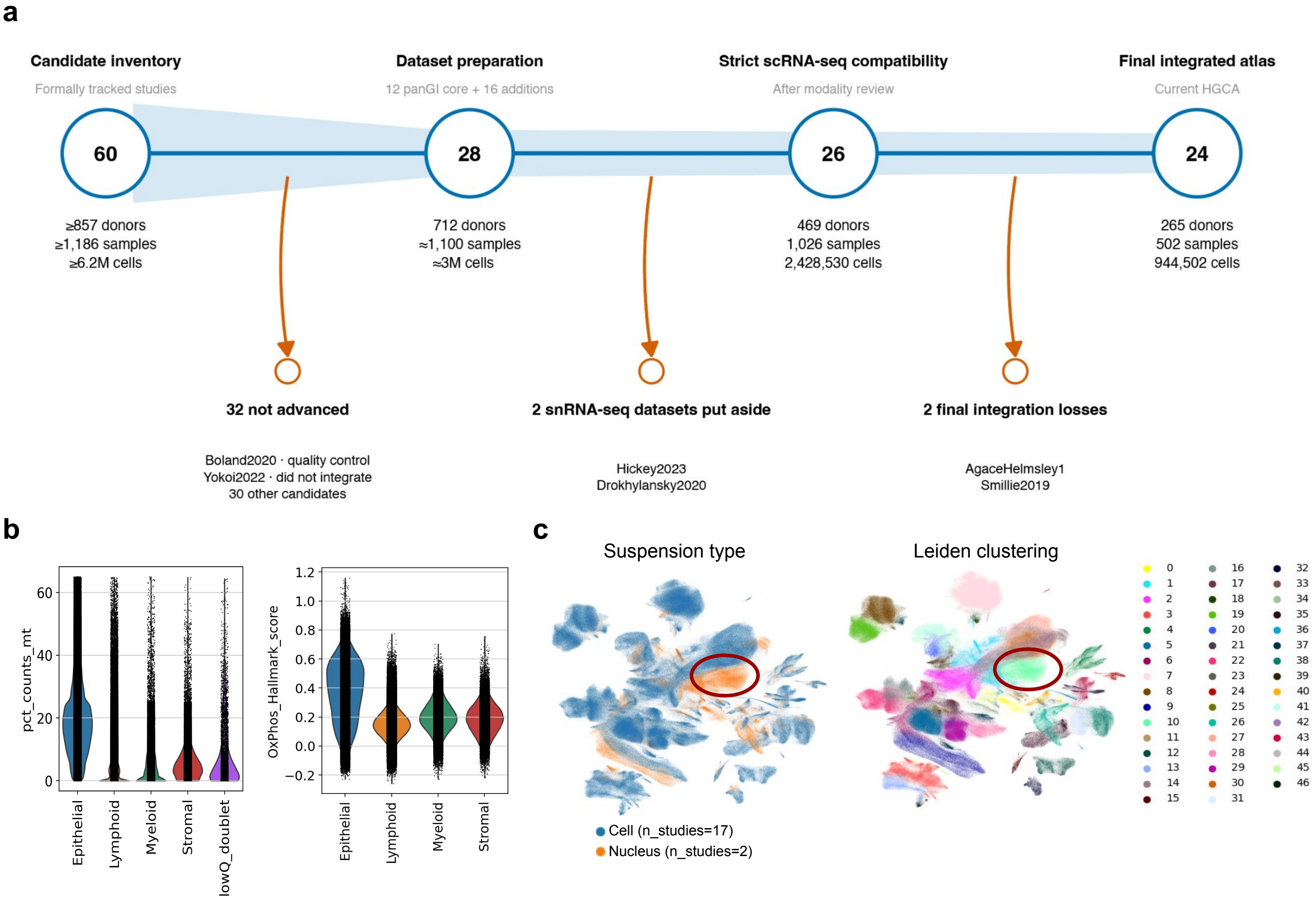


**Supplementary Figure 1 | Dataset selection, quality control and integration of datasets. a,** Overview of the dataset inclusion process. **b,** Lineage-specific distribution of the percentage of mitochondrial genes per cell (left) and respiratory activity quantified by gene set scoring using the MSigDB Hallmark gene set oxidative phosphorylation (OxPhos). The OxPhos gene set does not include any genes encoded by the mitochondrial genome and therefore supports that higher fractions of mitochondrial counts in epithelial cells is part of their cell identity and not a low quality signature. **c,** UMAP of an earlier benchmarking subset of the HGCA datasets visualizing the integrated embedding of single cell and single nucleus datasets. Leiden clustering driven by the suspension type and the low number of single nucleus datasets prevented robust evaluation of technical batch effects versus biologically-driven cell states.

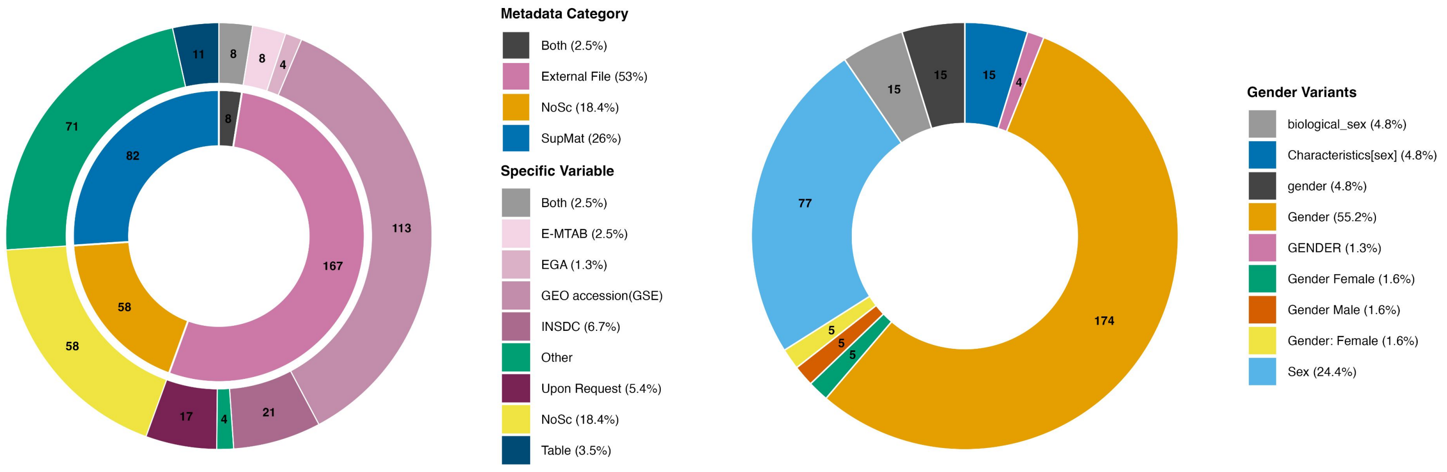


**Supplementary Figure 2 | Inconsistencies of metadata across single cell studies.** Availability, inconsistent data schema, and use of non-standardized terms make integrated analyses challenging.

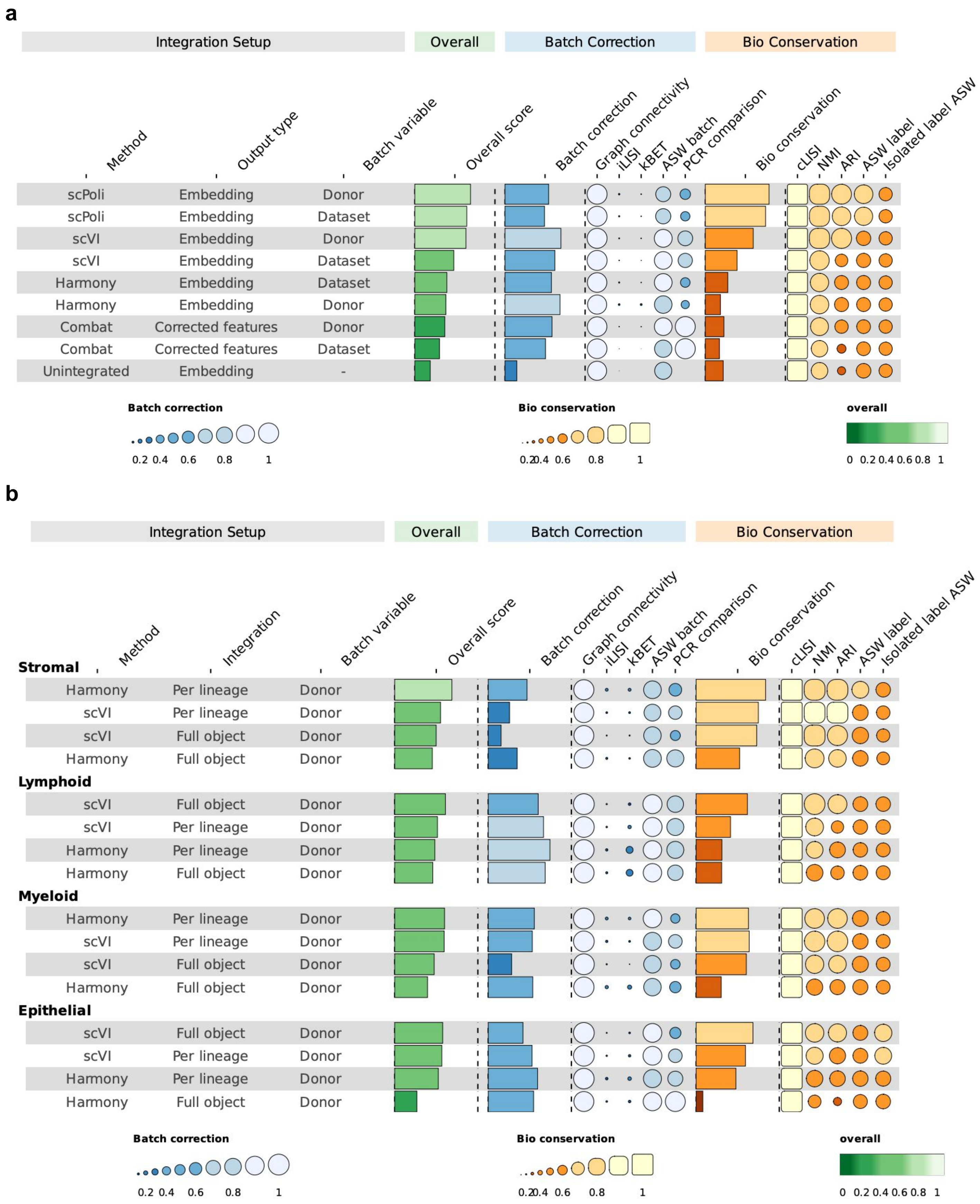


**Supplementary Figure 3 | Integration Benchmark using scAtlasTb.** Integration benchmarking on 50% of the samples in the HGCA balanced for dataset. Metric tables with 0-1 scaled scoring per metric and inverse for Bio conservation vs Batch correction such that a higher score means a better integration in both categories. **a,** First step integration benchmark on the full dataset across lineages using 3000 HVGs to select best performing method and batch covariate (scPoli not considered to be cell type label agnostic). **b,** Additional investigation of top performing methods on full versus lineage specific integrations. Better conservation of biological variation for scVI compared to Harmony in three of four lineages. Batch correction in scVI performs better in per-lineage integrations. For comparability, metrics on the full object were calculated based on lineage subsets of the full object including recalculation of the knn-graph per split.

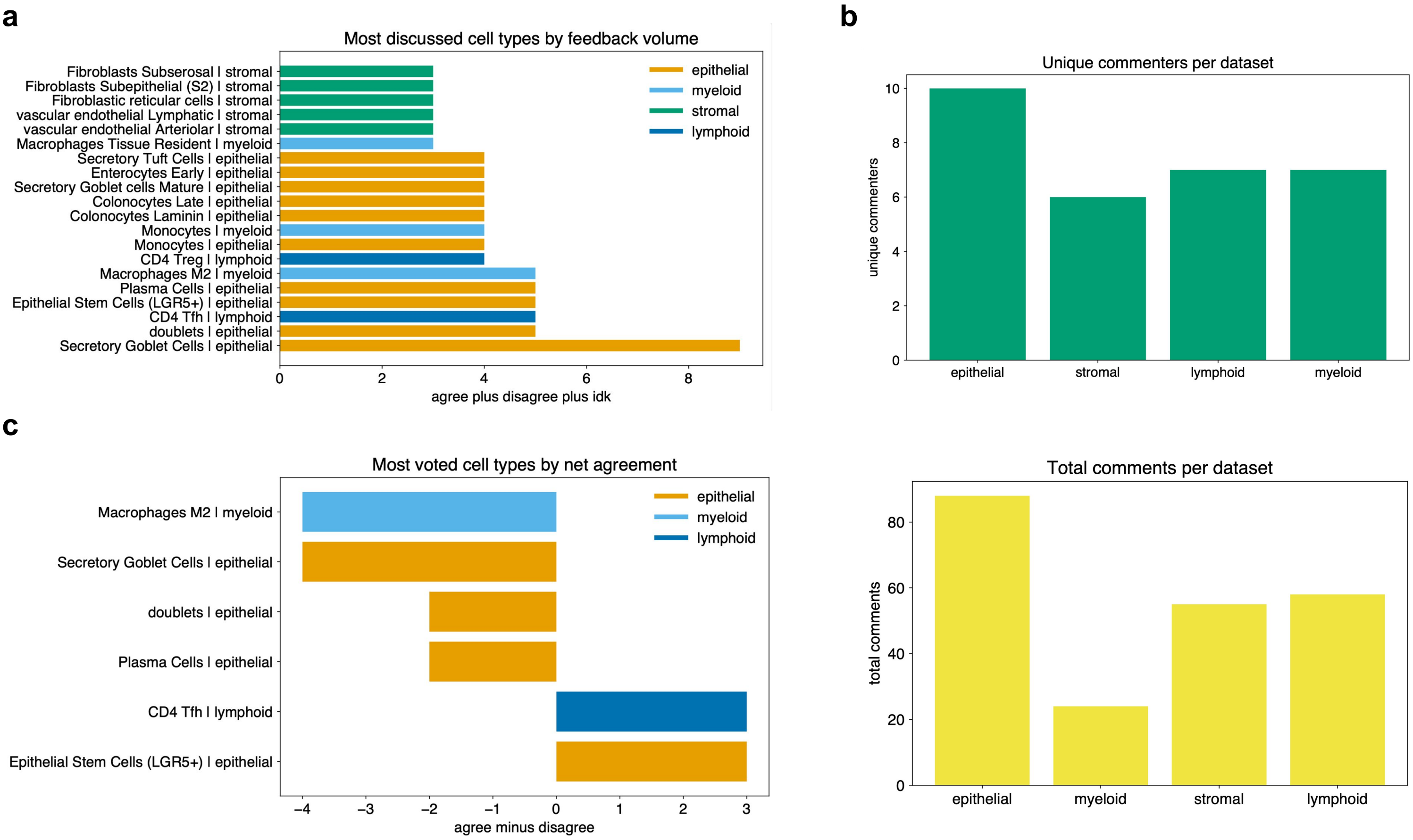


**Supplementary Figure 4 | CAP feedback on v0 labels. a,** Most voted-on v0 annotations across lineage objects, including "agree", "disagree", and "I don't know" (idk). The most common piece of feedback was to remove the word "Secretory" preceding the annotation "Secretory Goblet Cells" **b**, Unique commenters and total comments per lineage at the time of article submission. **c,** Most voted cell types arranged by net agreement, showing for which cell types there was the most consensus.

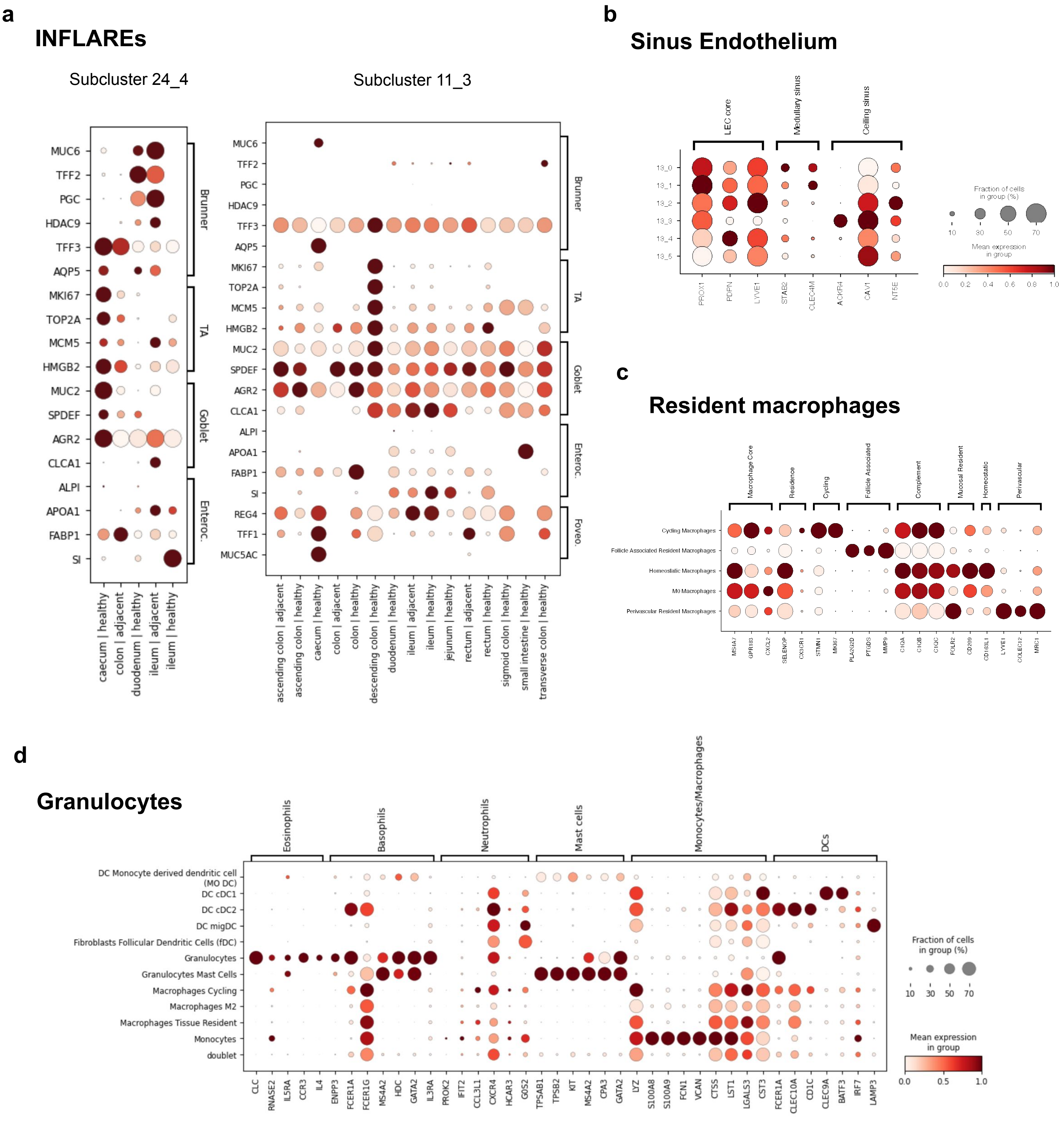


**Supplementary Figure 5 | Rare cell types.** **a,** Reidentification of INFLAREs as Brunner's gland and foveolar cells in integrated subclusters of the HGCA (MUC6 - Brunner's, MUC5AC - foveolar). **b**, Subtypes of sinus endothelial cells found by marker expression among subclusters of a lymphatic endothelial cluster in the stromal lineage. **c**, resident macrophage subtype markers by lineage-defining, shared program, and subtype specific markers. **d**, identification of rarely found granulocytes in the HGCA v0 label set as eosinophils by marker expression.

**a**

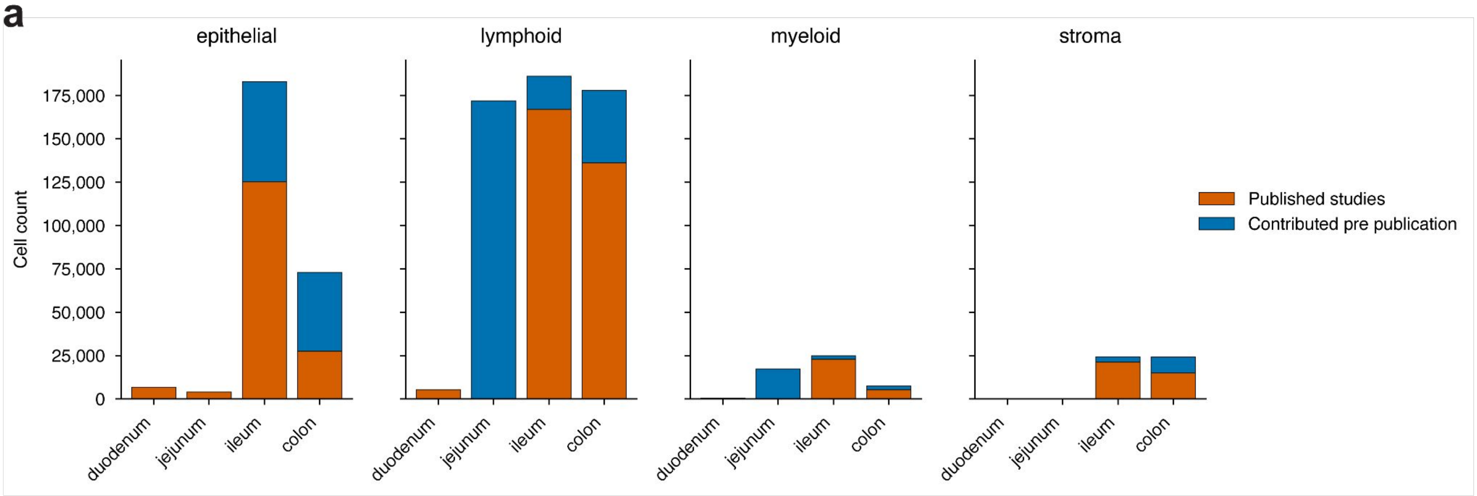


**Supplementary Figure 6 | Contribution of datasets pre-publication strengthened the atlas.**
**a,** Number of cells contributed by HGCA Bionetwork members per lineage and gut segment, colored by whether the dataset was contributed before or after publication and data release. **b,** Power analysis for differential expression (DE) between ileum and colon, showing increase in DE genes found once unpublished datasets are included. Samples included in the comparison are balanced according to the minimum number of them in either ileum or colon. DESeq2 formula included removal of dataset-specific batch effect.

**a**

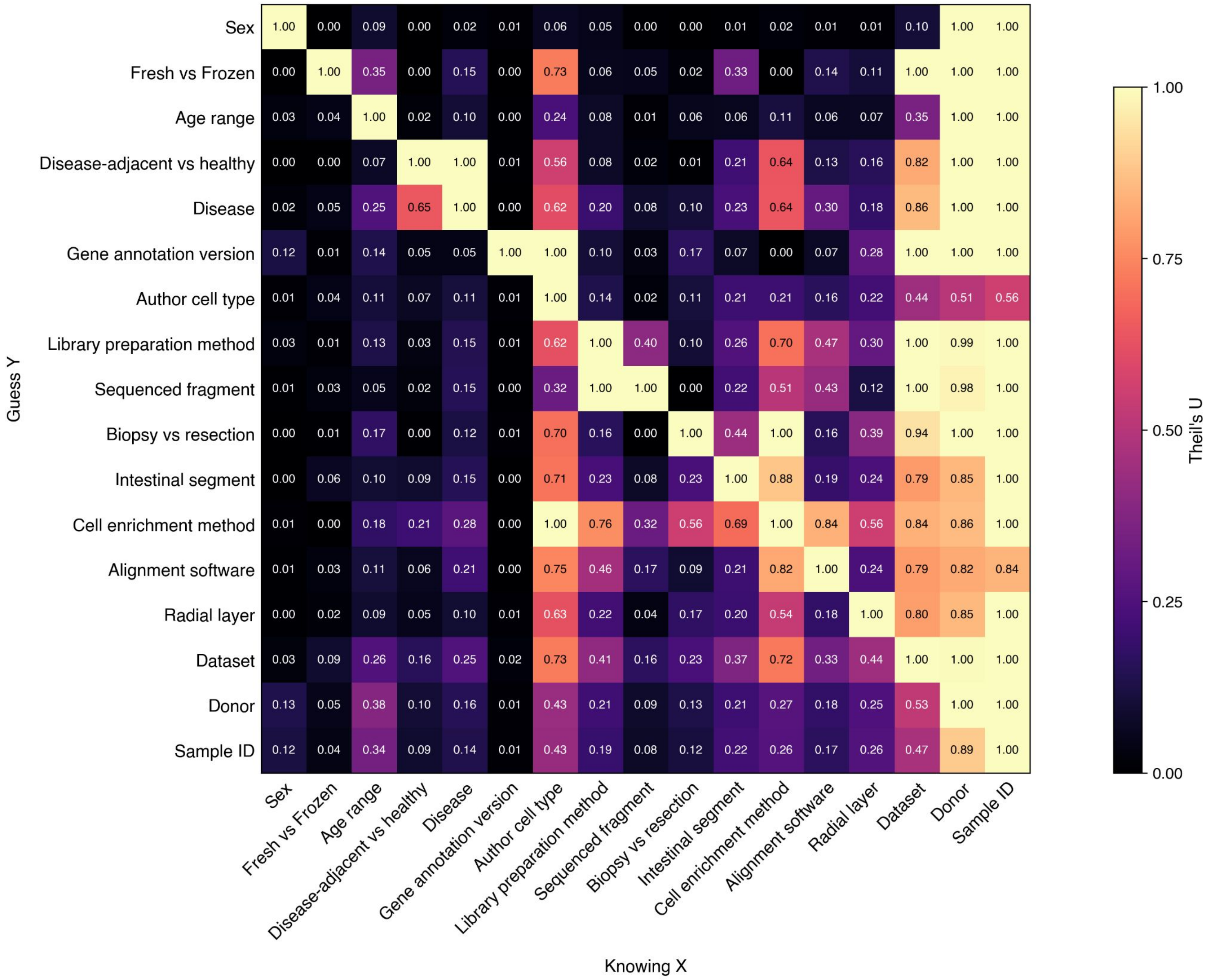


**Supplementary Figure 7 | Covariate confounding across the HGCA. a)** Theil's U heatmap showing variables confounded within and across datasets. Theil's U can be interpreted as normalized mutual information, and is a directional metric - using Dataset as an example, if you know which Sample a cell came from (last column of x axis), how well does that let you guess the other covariates.

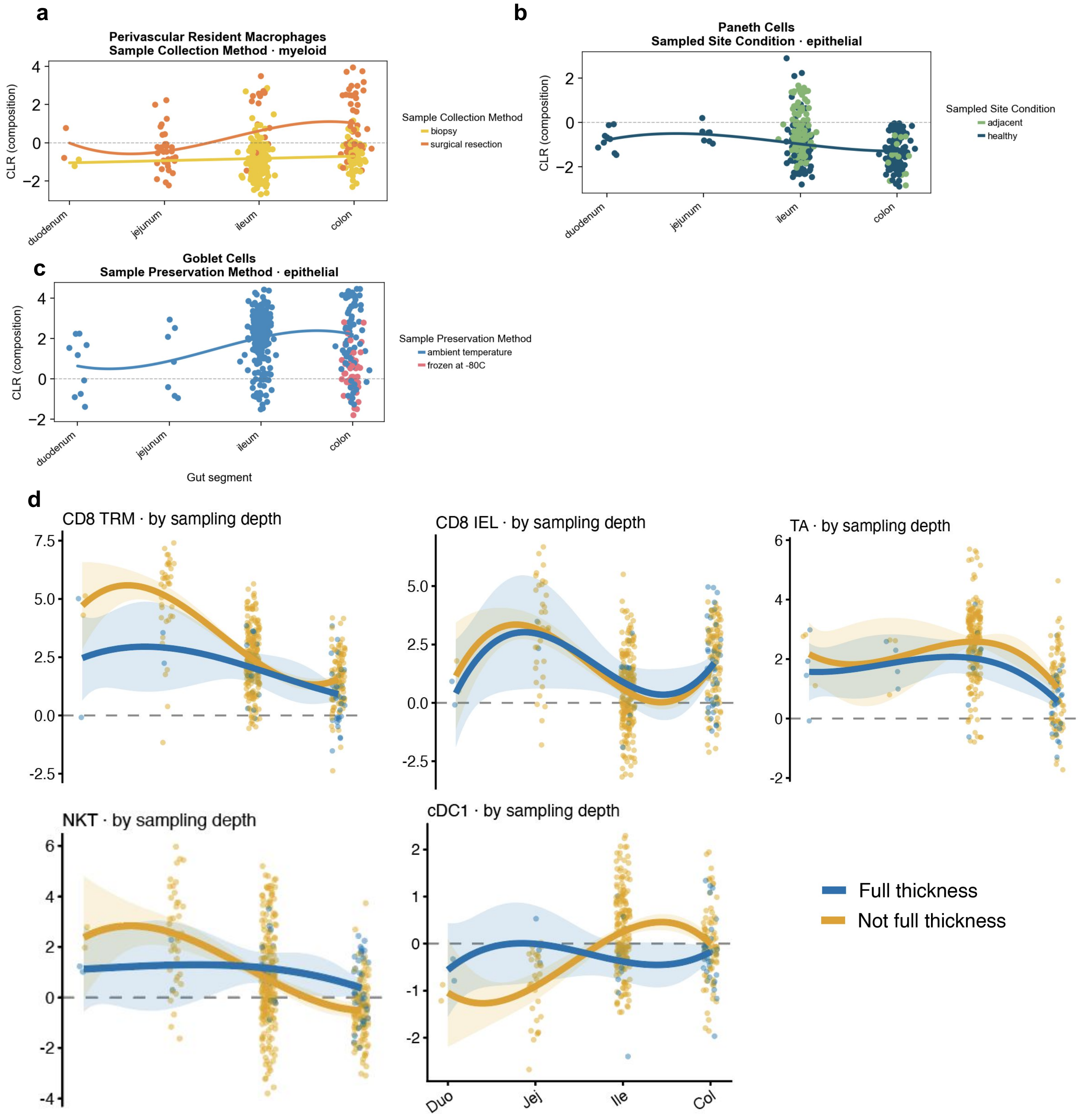


**Supplementary Figure 8 | Example compositions dependent on metadata covariates. a,** Perivascular resident macrophages are preferentially captured in surgical resections of the distal gut, **b,** Paneth cells are higher relative abundance in the uninvolved (disease-adjacent) Ileum of diseased patients, and **c,** Goblet cells are captured at lower rates if the protocol includes -80C freezing. **d,** CD8 TRM, CD8 IEL, TA, and NKT decrease in relative abundance along the segments of the gut, while cDC1s increase. These differences can depend on capture by full thickness vs radial layers.

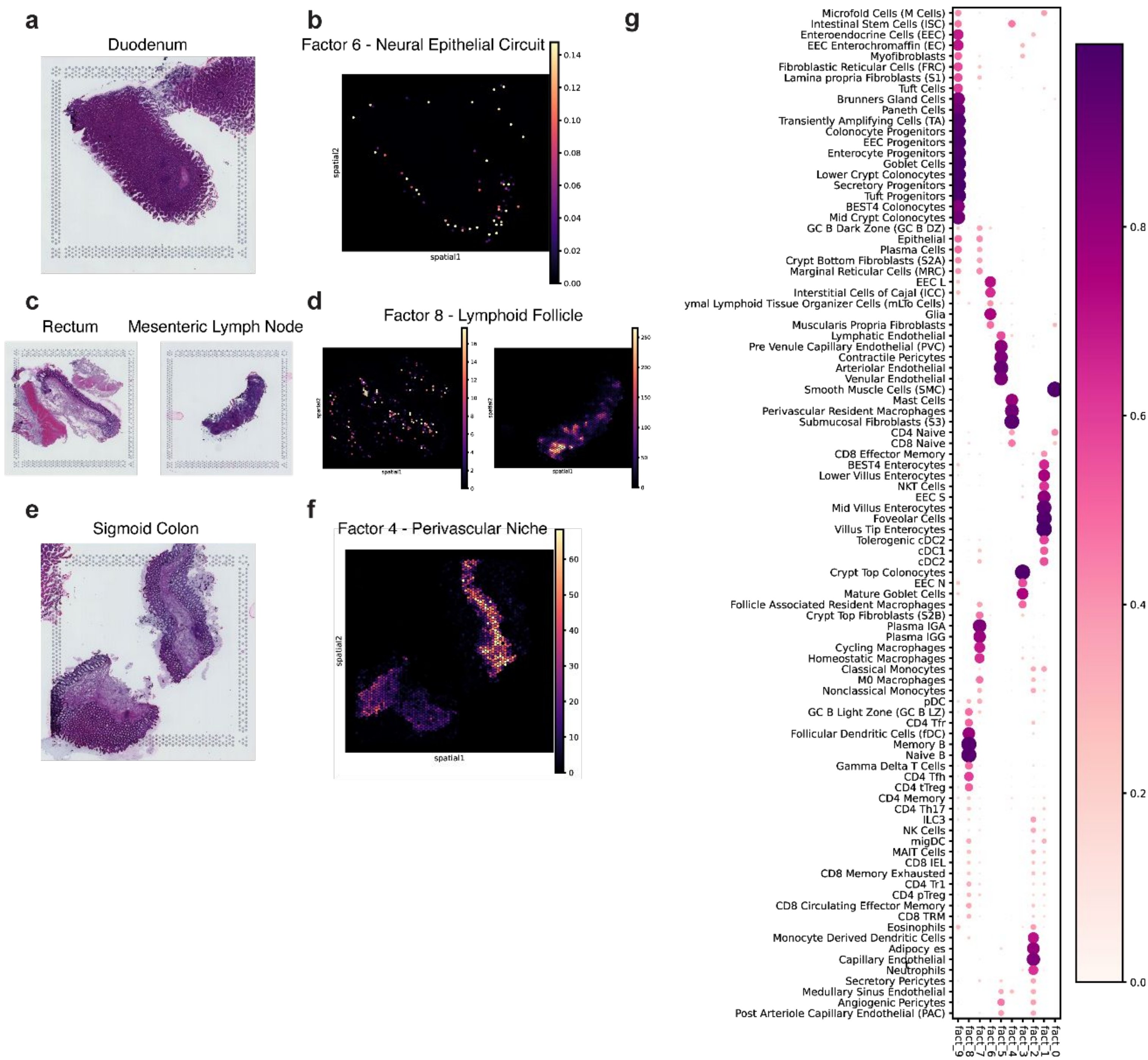

**Supplementary Figure 9 | Cell type localizations in 10X Visium. a,** Healthy duodenum H&E side by side with **b,** Cell2Location enrichment of neural epithelial circuit genes per Visium spot. **c,** Healthy rectum and mesenteric lymph node H&E images side by side with **d,** Cell2Location enrichment of a gene module containing lymphoid follicle niche cell types of these same H&E images. **e,** Healthy sigmoid colon H&E side by side with **f**, perivascular niche enrichment. See **g,** for factor (columns) by HGCA celltype (rows) loadings.

**a**

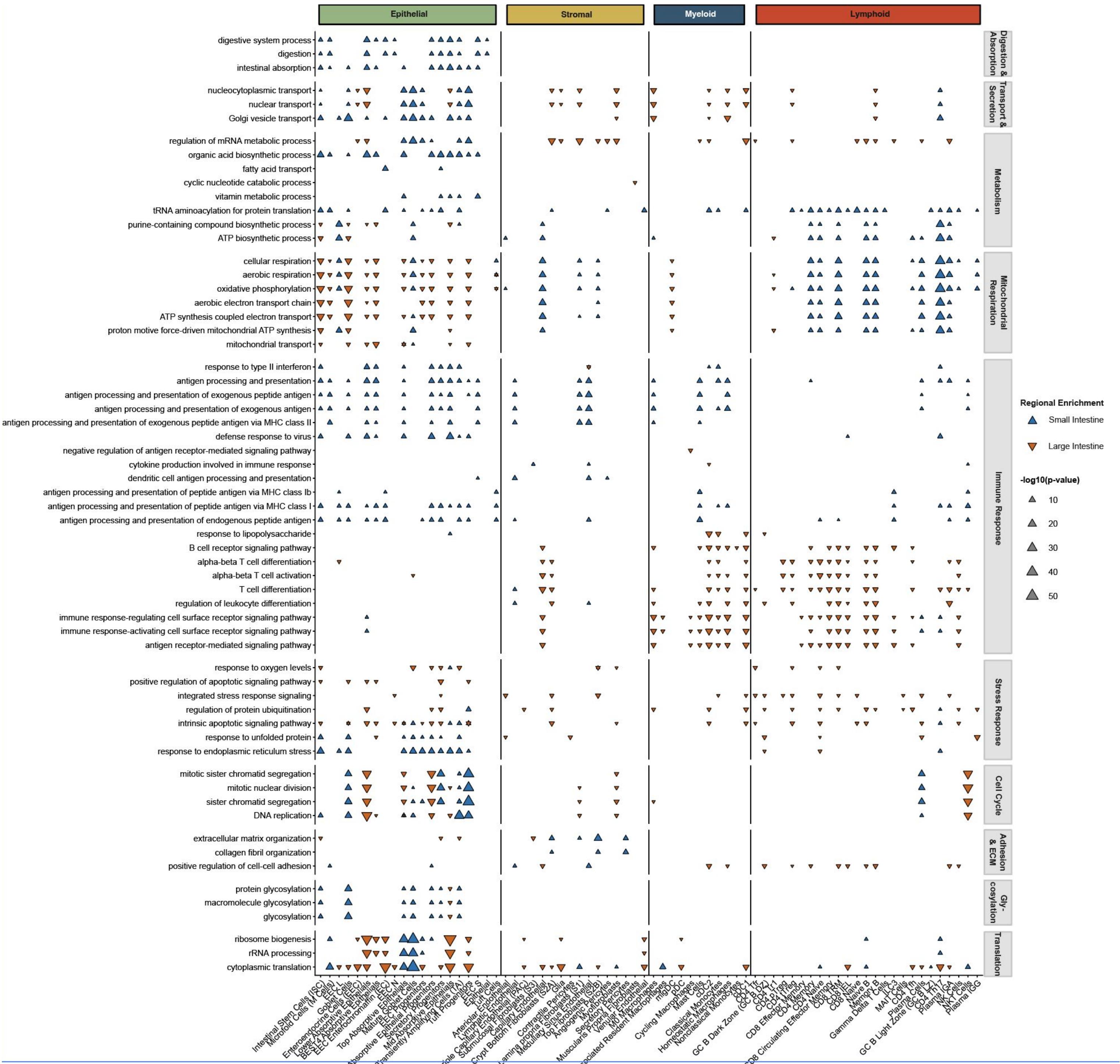


**b**

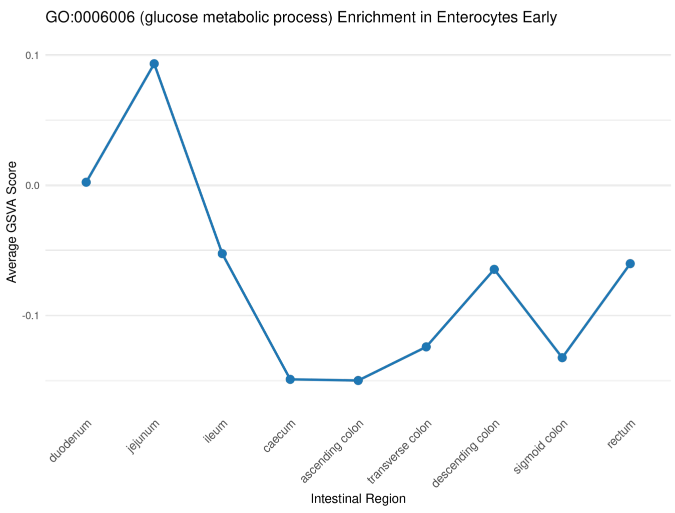


**Supplementary Figure 10 | Gene Ontology map of the major gut segments. a,** Triangles mark increased GSVA enrichment of GO term in **blue (upwards)** small intestine versus **orange (downwards)** large intestine, with triangle size determined by FDR-corrected p-values.

**a**

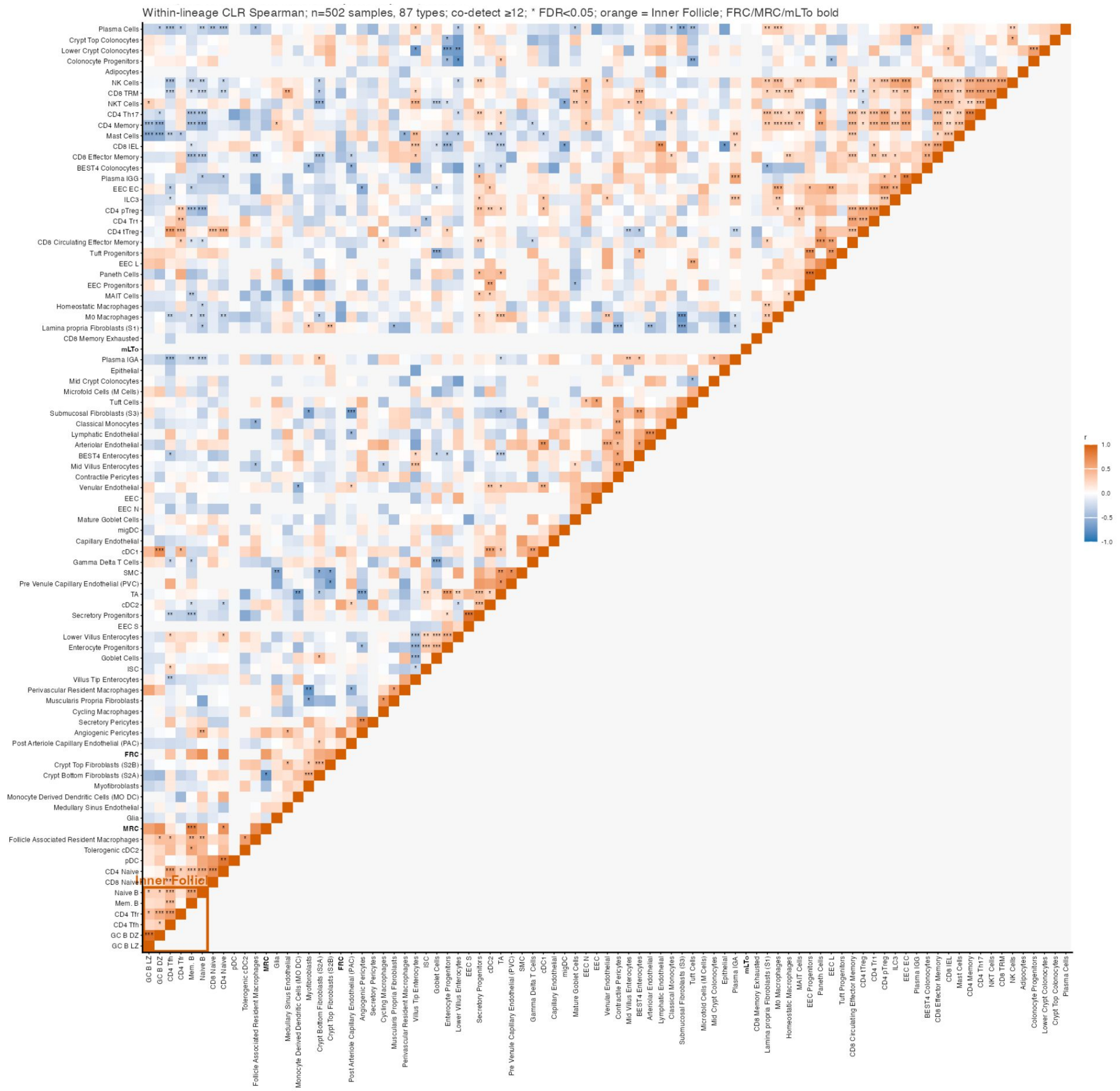


**Supplementary Figure 11 | Celltype compositions covarying across all samples. a,** Heatmap showing Spearman correlations between CLR transformed cell type compositions, with FDR adjusted p-values marked by stars: * p<0.05, ** p<0.01, *** p<0.005.

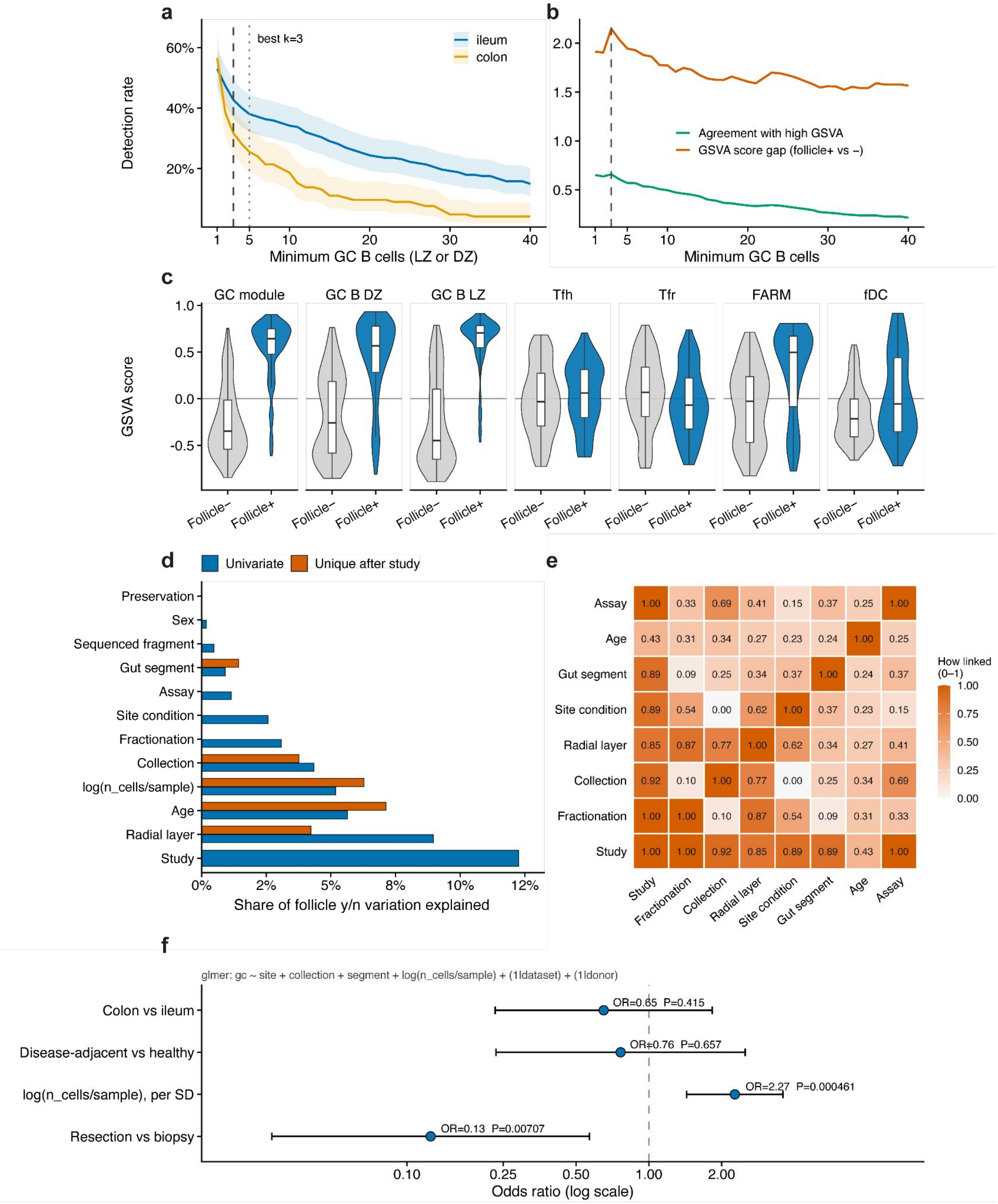


**Supplementary Figure 12 | Follicle presence can be estimated by a small GC B presence threshold.** **a,** Follicle detection rate versus k (GC B cutoff) (Wilson 95% CIs) for ileum and colon samples. Vertical lines mark potential cutoffs. **b**, How well each cutoff separates GC-module GSVA scores. Difference in mean GSVA scores between follicle+ and follicle- samples (orange), and agreement of the composition call with GSVA score > 0 (green). k=3 cutoff maximizes both heuristics. **c,** Sample-level GSVA for follicle-associated programs at k = 3, split by follicle- & follicle+. **d,** Share of follicle-call variation explained by each covariate ($R^2$ from logistic models) with univariate association (blue) and unique contribution after additive adjustment for study (orange). **e**, pairwise linkage among design covariates (Cramer's V; 0 = independent, 1 = perfectly associated), showing that chemical fractionation, collection method, and radial layer are together study nested, so that if chemical fractionation causes variation, it is confounded with these other covariates. **f,** Odds ratios from a binomial mixed model of follicle presence (k = 3 cutoff). Fractionation is not included due to **e.**

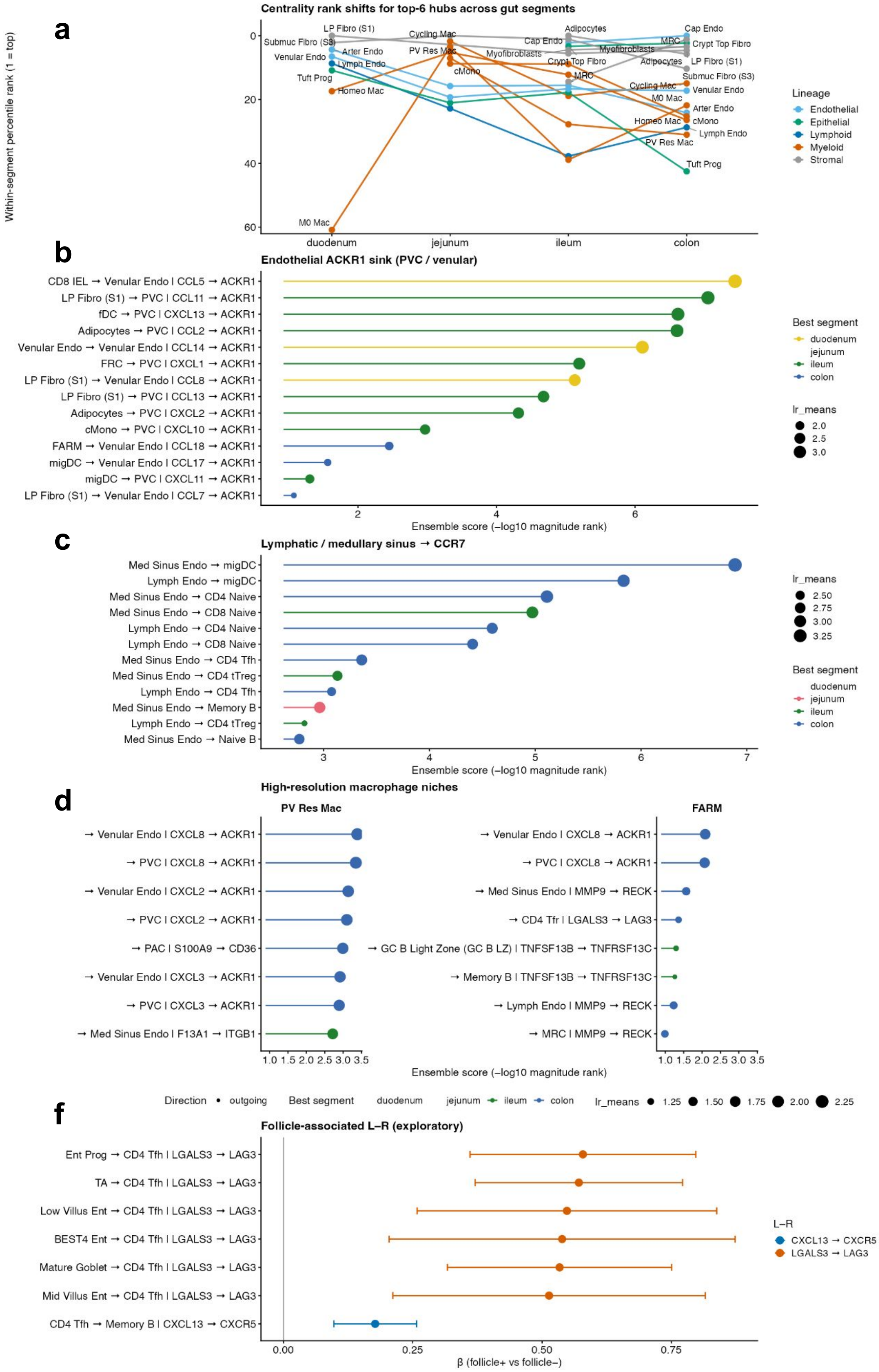


**Supplementary Figure 13 | Cell-cell communication across gut compartments with LIANA ensemble. a,** Top 6 cell types per gut segment are displayed across gut segments by LIANA ensemble of methods consensus degree centrality rank in CCC analysis. Lines connect each top ranked cell type across the gut segments. Lines are colored by cell type lineage. **b**, Top ranked interactions sending ligands to ACKR1 in Endothelial subtypes (PVC = pre venule capillary), colored by gut segment in which they occur **c**, Outgoing CCR7 signal from Lymphatic and sinus endothelial cells, showing top receiving cell types, colored by gut segment. Next displayed are top ranked outgoing signals excluding complement (C1Q*) from rare tissue resident macrophages: **d**, Perivascular resident macrophages (PV Res Mac) send CXCL2, CXCL3, and CXCL8 to ACKR1 on endothelial cells, while **e,** follicle associated resident macrophages (FARM) send similar signals alongside MMP9, TNFSF13B, and LGALS3 to follicle associated cell types. **f,** L-R CXCL13-CXCR5 and LGALS3-LAG3 L-R pairs were tested across cell types and associated with follicle presence in tissue samples with logistic models. L-R cell type interactions are ordered by logistic model beta and 95% confidence intervals across samples.

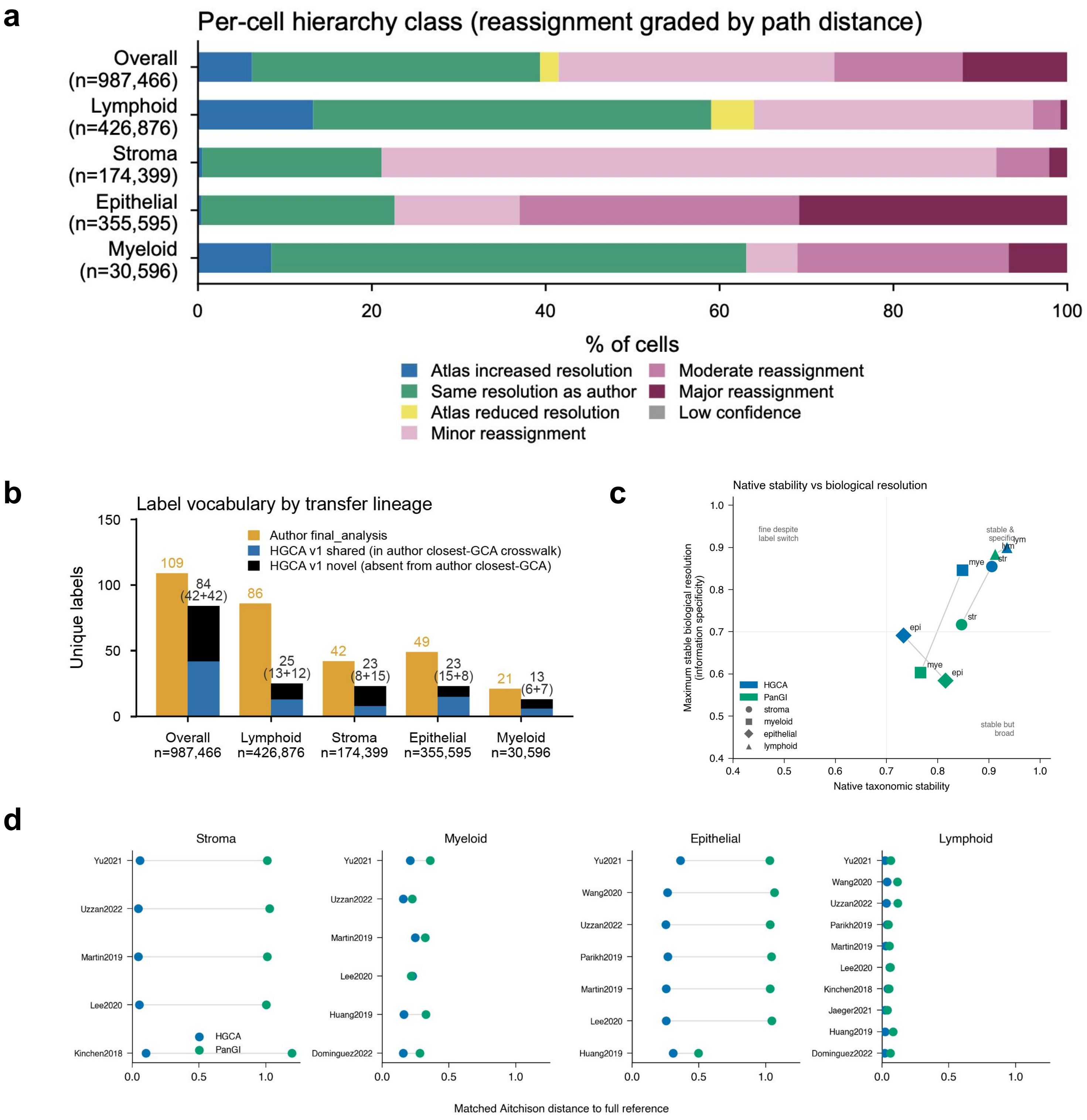


**Supplementary Figure 14 | Mapping resolution and stability for disease data depends on atlas constituent datasets. a,** Label set comparisons between TAURUS and HGCA, showing increase in number of healthy labels by reference mapping HGCA. **b,** Relationship between transferred HGCA identities and closest TAURUS author labels, classified per cell as increased, reduced, same resolution, or reassignment to another branch of the healthy cell-type hierarchy, with minor (within-level difference), moderate (two levels), and major (three or more levels) reassignments. **c,** Native label set mapping stability (x axis) against maximum stable biological resolution (y axis) for each lineage model in the shared LODO jackknife test, showing epithelial labels are more stable in PanGI because they map a lower maximum resolution, while HGCA myeloid and stromal lineages are both higher resolution and more stable across identities. **d**, Differences in cell type composition by Aitchison Distance (CLR transformed Euclidean distance) after one study was removed from each reference atlas. Distances were calculated after matching HGCA and PanGI to the finest common denominator label.

a

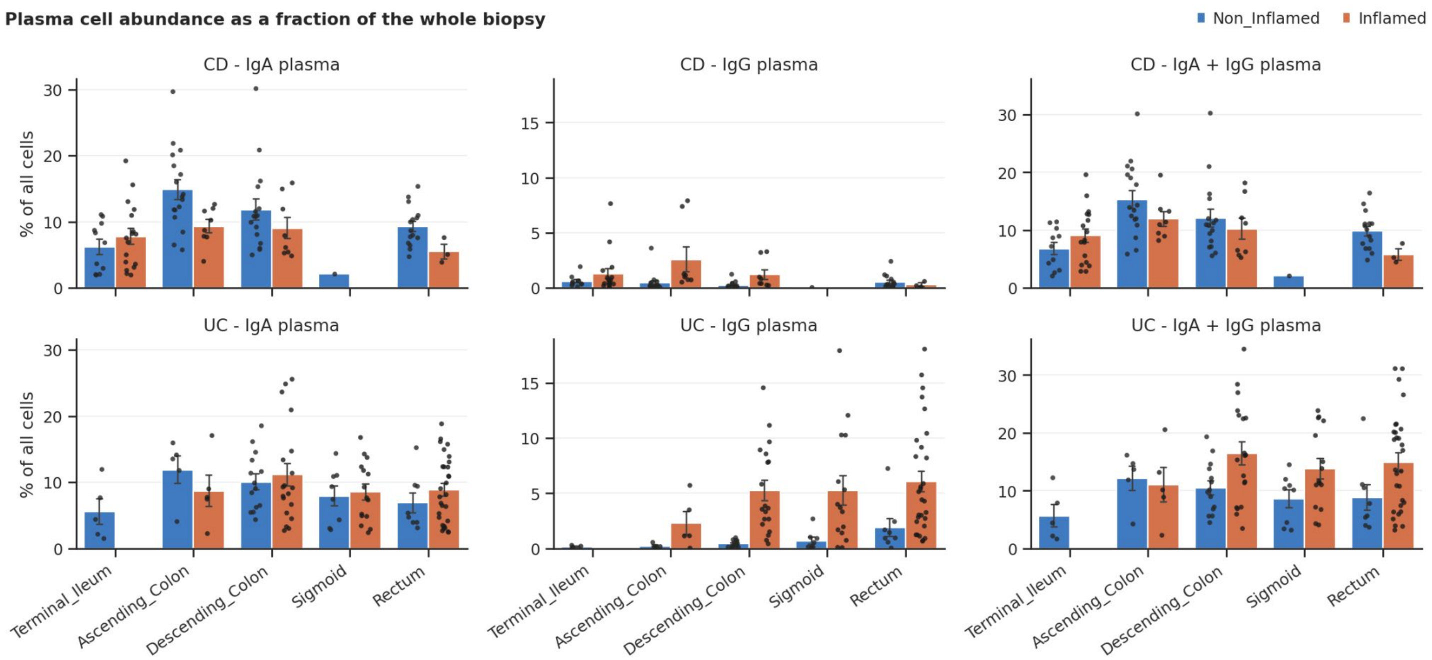


b

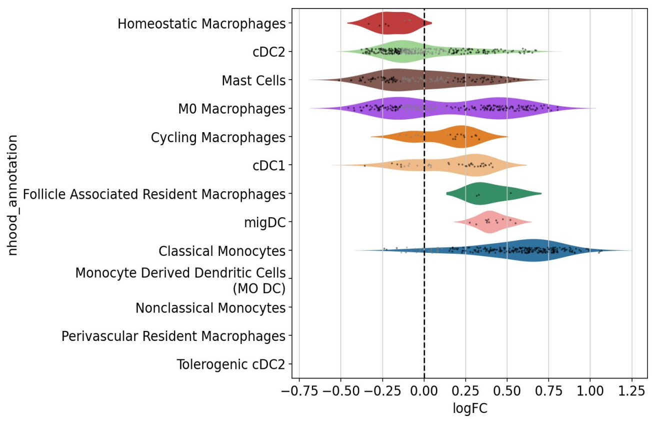


**Supplementary Figure 15 | Detailed mapping results of TAURUS cohort to the HGCA and inflammation signatures. a,** Heatmap showing original author annotation of the TAURUS story and predicted labels from HGCA label transfer. **b,** IGG /IGA plasma cell abundance in CD and UC across anatomic locations. **c,** Milo results in the Myeloid compartment with inflammation score as the variable for identifying enriched neighborhoods

**a**

| | | Subtotals | | Biological | | | | | | | | Technical | | |
|---|---|---|---|---|---|---|---|---|---|---|---|---|---|---|
| representation | Total | Biological | Technical | Severity | Remission | Inflammation | Disease | Treatment | Age | Gender | Disease Duration | Batch | Library type | n cells |
| GloScope | 0.52 | 0.46 | 0.63 | 0.56 | 0.5 | 0.66 | 0.44 | 0.31 | 0.54 | 0.31 | 0.39 | 0.7 | 0.66 | 0.52 |
| CT pseudobulk | 0.47 | 0.45 | 0.5 | 0.56 | 0.38 | 0.56 | 0.29 | 0.4 | 0.52 | 0.43 | 0.47 | 0.68 | 0.29 | 0.52 |
| composition (CLR | 0.44 | 0.34 | 0.64 | 0.27 | 0.35 | 0.43 | 0.19 | 0.4 | 0.35 | 0.44 | 0.33 | 0.76 | 0.57 | 0.58 |
| MOFA | 0.42 | 0.32 | 0.61 | 0.42 | 0.23 | 0.38 | 0.0 | 0.48 | 0.29 | 0.33 | 0.46 | 0.71 | 0.53 | 0.59 |
| PILOT | 0.41 | 0.21 | 0.81 | 0.41 | 0.34 | 0.22 | 0.0 | 0.2 | 0.22 | 0.12 | 0.17 | 0.91 | 0.93 | 0.59 |
| Pseudobulk | 0.38 | 0.2 | 0.74 | 0.22 | 0.23 | 0.33 | 0.0 | 0.36 | 0.06 | 0.16 | 0.19 | 0.82 | 0.61 | 0.78 |
| Random vector | 0.35 | 0.02 | 1.0 | 0.0 | 0.02 | 0.01 | 0.0 | 0.0 | 0.13 | 0.0 | 0.0 | 1.0 | 1.0 | 1.0 |
| MixMIL | 0.32 | 0.09 | 0.77 | 0.02 | 0.0 | 0.18 | 0.0 | 0.04 | 0.25 | 0.21 | 0.0 | 0.96 | 0.8 | 0.55 |

**b** **c**

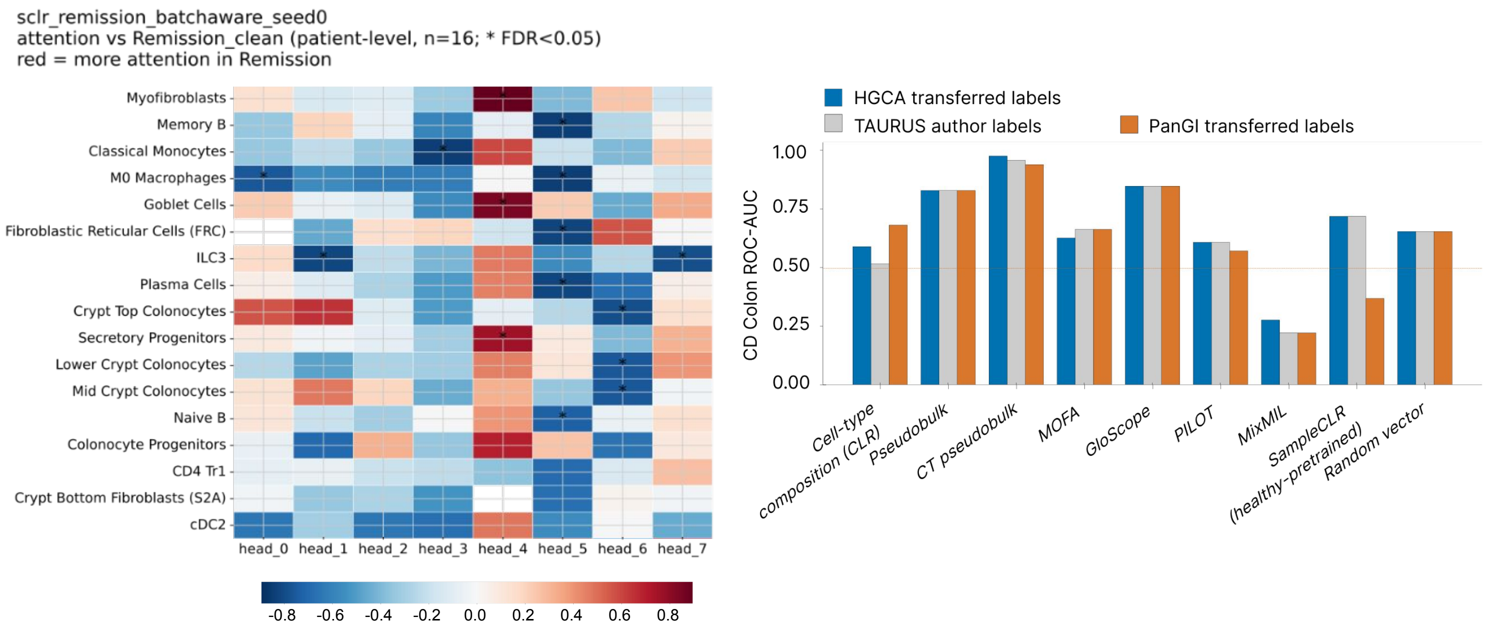


**Supplementary Figure 16 | Taurus Crohn's Colon patpy sample embedding benchmark. a,** Patpy benchmarks of top performing embeddings for Taurus CD Colon samples using HGCA labels, showing that GloScope, a label-free method, performs best across biological signal preservation and technical batch removal. **b**, Spearman correlation of cell types with attention head weights in SampleCLR embedding trained on pre-treatment Crohn's colon samples from TAURUS, with learning targeted at future remission status. **c,** ROC-AUC for predicting eventual anti-TNF remission in pretreatment colonic CD samples across sample-representation methods using HGCA-transferred, TAURUS author, or PanGI-transferred cell type labels. Horizontal line indicates exact randomness and random-vector representation provides an empirical baseline.

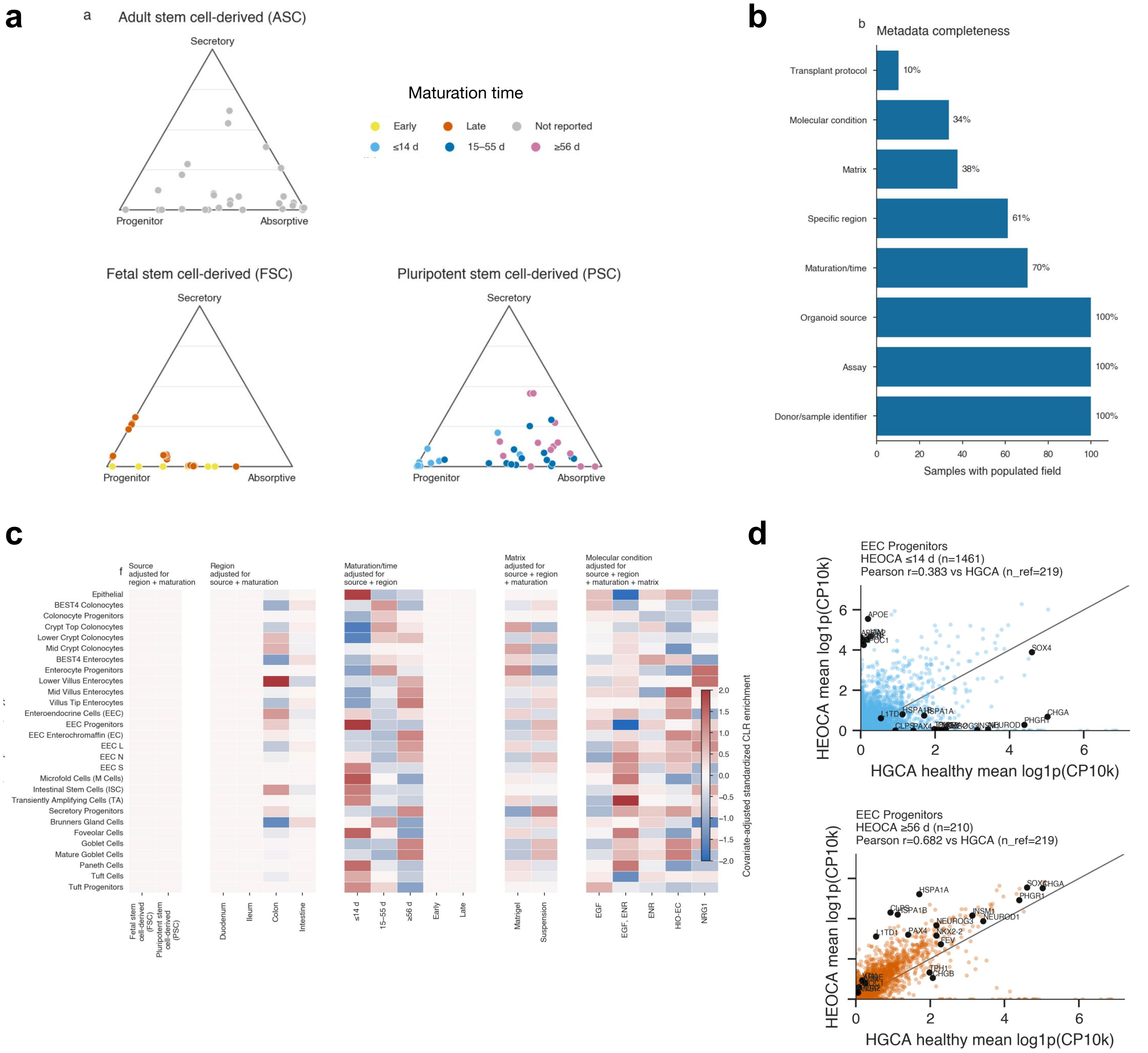


**Supplementary Figure 17 | Published HEOCA gut organoids differ in epithelial branch allocation and adult-reference correspondence, while protocol metadata remain incomplete**

**a,** Sample epithelial compositions shown as mutually exclusive progenitor, differentiated absorptive and differentiated secretory fractions, faceted by adult stem cell-derived (ASC), fetal stem cell-derived (FSC) and pluripotent stem cell-derived (PSC) organoids. Points represent samples and colours denote reported maturation/time. BEST4 identities are classified as absorptive. **b,** Percentage of 98 samples with each metadata field populated. Completeness indicates presence, not harmonization or independent experimental variation. **c,** HGCA epithelial-subtype enrichment across source, region, maturation, matrix and molecular-condition categories. Colour denotes standardized CLR enrichment after adjustment for the covariates indicated above each heatmap; blank cells were not estimable. Effects remain descriptive and may retain publication confounding. **d,** EEC progenitors begin to express lineage-defining markers only after maturation *in vitro*
