## Supplementary material for "The Human Gut Cell Atlas v1.0: A Unified, Community-Built Reference of Cellular Diversity Across the Gut": HGCA Gut Bionetwork contributors

All contributors listed below are also named individually in the manuscript byline.

Candace M. Cham[1], Eugene B. Chang[2], Can Ergen[3], Donna L. Farber[4,5], Mennatallah Ghouraba[6], Jason Koval[2], Christopher J. Lengner[7,8], Michal Mark[9], David T. Rubin[10], Peter A. Sims[11,12], Steven B. Wells[11,4], Bingqing Xie[13], Nir Yosef[9,14], Yusen Zhou[15], David Fawkner-Corbett[16,17], Thomas M. Fenton[18], Scott T. Magness[19,20], Shalev Itzkovitz[21], Monika Krzak[22], Rebecca E McIntyre[23], Adrienne Luoma[24], Gregory J. Wickham[25], Joseph Burclaff[26], Shengbao Suo[27], Liza Konnikova[28], Keith Alexander Breau[29], Weihong Gu[30], Espen S Baekkevold[31,32]

1 Section of Gastroenterology, Hepatology, and Nutrition, Department of Medicine, The University of Chicago, Chicago, IL, USA.

2 UChicago Medicine, Chicago, IL, USA.

3 Wurzburg Institute of Systems Immunology, Julius-Maximilians-Universitat Wurzburg, Wurzburg, Germany.

4 Department of Microbiology and Immunology, Columbia University Irving Medical Center, New York, NY, USA.

5 Department of Surgery, Columbia University Irving Medical Center, New York, NY, USA.

6 Barts Cancer Institute, Queen Mary University of London, London, United Kingdom.

7 Department of Biomedical Sciences, School of Veterinary Medicine, University of Pennsylvania, Philadelphia, Pennsylvania, USA.

8 Institute for Regenerative Medicine, University of Pennsylvania, Philadelphia, Pennsylvania, USA.

9 Department of Systems Immunology, Weizmann Institute, Rehovot, Israel.

10 Department of Medicine, The University of Chicago, Chicago, IL, USA.

11 Department of Systems Biology, Columbia University Irving Medical Center, New York, NY, USA.

12 Department of Biochemistry & Molecular Biophysics, Columbia University Irving Medical Center, New York, NY, USA.

13 Department of Internal Medicine, UT Southwestern Medical Center, Dallas, TX, USA.

14 Department of Electrical Engineering and Computer Science and Center for Computational Biology, University of California, Berkeley, CA, USA.

15 Division of Gastroenterology, Hepatology, and Nutrition, Department of Pediatrics, University of Pennsylvania Perelman School of Medicine and Children's Hospital of Philadelphia, Philadelphia, PA, USA.

16 Medical Research Council Translational Immune Discovery Unit, Weatherall Institute of Molecular Medicine, University of Oxford, Oxford, United Kingdom.

17 Academic Paediatric Surgery Unit, Nuffield Department of Surgical Sciences, University of Oxford, Oxford, United Kingdom.

18 Technical University of Denmark, Denmark.

19 UNC/NCSU Lampe Joint Departments of Biomedical Engineering, University of North Carolina at Chapel Hill and North Carolina State University, NC, USA.

20 Departments of Medicine, Cell Biology and Physiology, University of North Carolina at Chapel Hill, Chapel Hill, NC, USA.

21 Department of Molecular Cell Biology, Weizmann Institute of Science, Rehovot, Israel.

22 Wellcome Sanger Institute. Hinxton, Cambridgeshire, CB10 1SA, United Kingdom.

23 Relation Therapeutics, London, United Kingdom.

24 Dana-Farber Cancer Institute, Boston, MA, USA.

25 Earlham Institute, Norwich Research Park, Norwich, NR4 7UG, United Kingdom.

26 Department of Molecular Biomedical Sciences, College of Veterinary Medicine, North Carolina State University, Raleigh, NC 27603, USA.

27 Guangzhou National Laboratory, Guangzhou, China.

28 Departments of Pediatrics, Immunobiology and OB, GYN and Reproductive Sciences, Yale University, New Haven, CT, USA.

29 University of North Carolina at Chapel Hill, Chapel Hill, NC, USA.

30 Yale School of Medicine, New Haven, CT, USA.

31 Department of Pathology, Oslo University Hospital, Oslo, Norway.

32 Institute of Oral Biology, University of Oslo, Oslo, Norway.

Present address: Monika Krzak, Institute of Metabolic Science, Department of Clinical Biochemistry, University of Cambridge, Cambridge, United Kingdom.
